# Chemical crosslinking enables profiling of endogenous cell surface galectin-3 interactions

**DOI:** 10.64898/2026.11.06.764912

**Authors:** Blake W Krisko, Aswath Balakrishnan, Andrew Wheat, Christofer D Sánchez, Clinton Yu, Ria Bhakta, Luna Witchey, Minas Minasyan, Lan Huang, Haik Mkhikian

## Abstract

Galectin-3 is a β-galactoside-binding lectin that organizes glycoproteins into cell surface lattices (multivalent interaction networks) that regulate biology spanning immune function, fibrosis, heart disease, and cancer progression. Galectin-3 engages promiscuous but low-affinity glycan ligands, posing technical challenges for faithfully mapping the cell surface galectin-3 interactome. Here, we introduce *Crosslinking-assisted Galectin Lattice Affinity Purification Mass Spectrometry* (xGL-APMS), a facile and stringent proteome-scale approach that captures galectin-3 interactions natively at the cell surface, without the need for exogenous galectin-3, and is extendable to XL-MS. We identified 125 N-glycan branching-dependent surface galectin-3 interactors on Jurkat cells, validated interactions with six orthogonal approaches, and compared low and high-dose galectin-3 interactomes, revealing differences in both distribution and interactome composition. Finally, without exogenous or modified bait, we performed immediate crosslinking of endogenous galectin-3 to identify 69 galectin-3-associated proteins on primary mouse thymocytes. Together, these approaches provide a readily adoptable framework for measuring endogenous surface galectin-3 and profiling its interacting partners.

## Introduction

The T cell surface is a dynamic molecular landscape that coordinates immune biology. Asparagine (N)-linked glycosylation, found on nearly all cell surface proteins, is an intricate sub-landscape governing protein folding, signaling activity, cell surface retention and carbohydrate-mediated protein interactions^1–7^. During N-glycan maturation, the Golgi-resident enzymes N-acetylglucosaminyltransferase I, II, IV, and V (encoded by *MGAT1*, *MGAT2*, *MGAT4a/b*, and *MGAT5*) add N-acetylglucosamine (GlcNAc) branches to generate mono-, bi-, tri-, and tetra-antennary N-glycan structures (Fig. S1a). Galactosylation of these branches generates N-acetyllactosamine (LacNAc), the major T cell ligand for a family of β-galactoside-binding proteins known as galectins^8–10^.

Galectin-3, one of 15 mammalian galectins, preferentially binds highly branched N-glycoforms to form multivalent lattices that regulate both T cell receptor activation and surface protein retention^11–14^ (Fig. S1b). Accordingly, N-glycan branching and galectin-3 have an increasingly appreciated role in conditions spanning autoimmunity, immune aging, cancer, heart disease, and fibrosis^15–21^. Yet, despite this growing disease relevance and clinical interest, proteome-wide characterization of native and endogenous galectin-3 lattices remains challenging. Mapping surface galectin-3 networks could clarify how galectin-3 contributes to disease and inform galectin-3 directed interventions.

Sensitive and accurate mapping of galectin-3 networks is hindered by several challenges. Traditional protein-protein interaction approaches, such as affinity purification of cell lysates on immobilized bait columns, remain a widely used means of proteomically profiling galectin-3^22,23^. However, lysate-based approaches do not reproduce the cell surface architecture under which physiological galectin-3 interactions form, returning sets of proteins with concerns of false positive and false negative discovery. The development of galectin-3 APEX2 proximity labeling enabled mapping of galectin-3 interactomes natively at the cell surface^24,25^. Although an important advance, notable limitations of this approach include: (i) fusion of a peroxidase domain, which may structurally or functionally perturb galectin-3, (ii) labeling occurs within a ∼20 nm radius, reporting the “protein neighborhood” rather than physical interactors, (iii) broad adoption or extension to other lectins requires bespoke fusion protein production and validation, and critically, (iv) the fusion protein must be overexpressed or added exogenously, precluding direct extension to endogenous and *in vivo* systems^26,27^. Metabolic glycan labeling has also been applied to probe surface interactions in live cells. In these approaches, modified monosaccharides supplied in culture are incorporated into glycans and then used to map glycan-protein interactions^28–30^. However, labeling is restricted to those monosaccharides which can tolerate modification (e.g., sialic acid), is biased toward glycoforms into which the modified sugars incorporate, and the artificial sugars themselves may perturb physiological interactions.

Direct chemical crosslinking is widely used to capture transient biomolecular interactions. For instance, formaldehyde crosslinking and purification enables profiling of transcription factor binding sites on DNA (ChIP-seq)^31,32^. For protein-protein interactions, amino acid side-chain-reactive linkers enable preservation of complex integrity through sample preparation and limit sample loss^33–36^. Crosslinking-based analyses have yielded deep protein-DNA and protein-protein interaction profiles across the nucleus, mitochondria, cytosol, and in limited capacity the cell surface^37–40^.

Here, we leverage chemical crosslinking to develop Crosslinking-assisted Galectin Lattice Affinity Purification Mass Spectrometry (xGL-APMS). xGL-APMS improves sensitivity for detecting surface galectin-3, enables high-confidence identification of N-glycan branching-dependent galectin-3 interactions, and identifies the endogenous galectin-3 interactome on primary mouse thymocytes. Together, immediate crosslinking and xGL-APMS provide high-resolution insight into galectin-3-mediated immunology and open a path to interrogating galectin-glycoprotein dynamics without requiring galectin/glycan modification or supplementation with exogenous proteins.

## Results

### Immediate crosslinking stabilizes glycan-dependent surface galectin-3

N-glycan branching is dynamically regulated during T cell development, decreasing sharply as CD4⁺CD8⁺ double positive (DP) thymocytes mature into CD4⁺ and CD8⁺ single positive (SP) thymocytes^41^. We initially asked whether this shift produced corresponding changes in endogenous surface galectin-3. Using flow cytometry, however, wild-type C57BL/6J DP and SP thymocytes displayed no galectin-3 signal above *Lgals3^-/-^* controls by median fluorescence intensity (MFI) despite established galectin-3 expression (Fig. 1a, Fig. S1c-e)^42^. Incubation with recombinant galectin-3 (rGAL-3) produced a modest increase in signal, confirming that galectin-3 is detectable in principle (Fig. 1a, Fig. S1e). Consistent with the inability to co-purify TCR and PTPRC (CD45) with galectin-3 without stabilization^12,43,44^, we suspected that our poor galectin-3 detection by flow cytometry reflected loss of surface galectin-3 rather than a low ceiling on surface occupancy.

**FIGURE 1:**
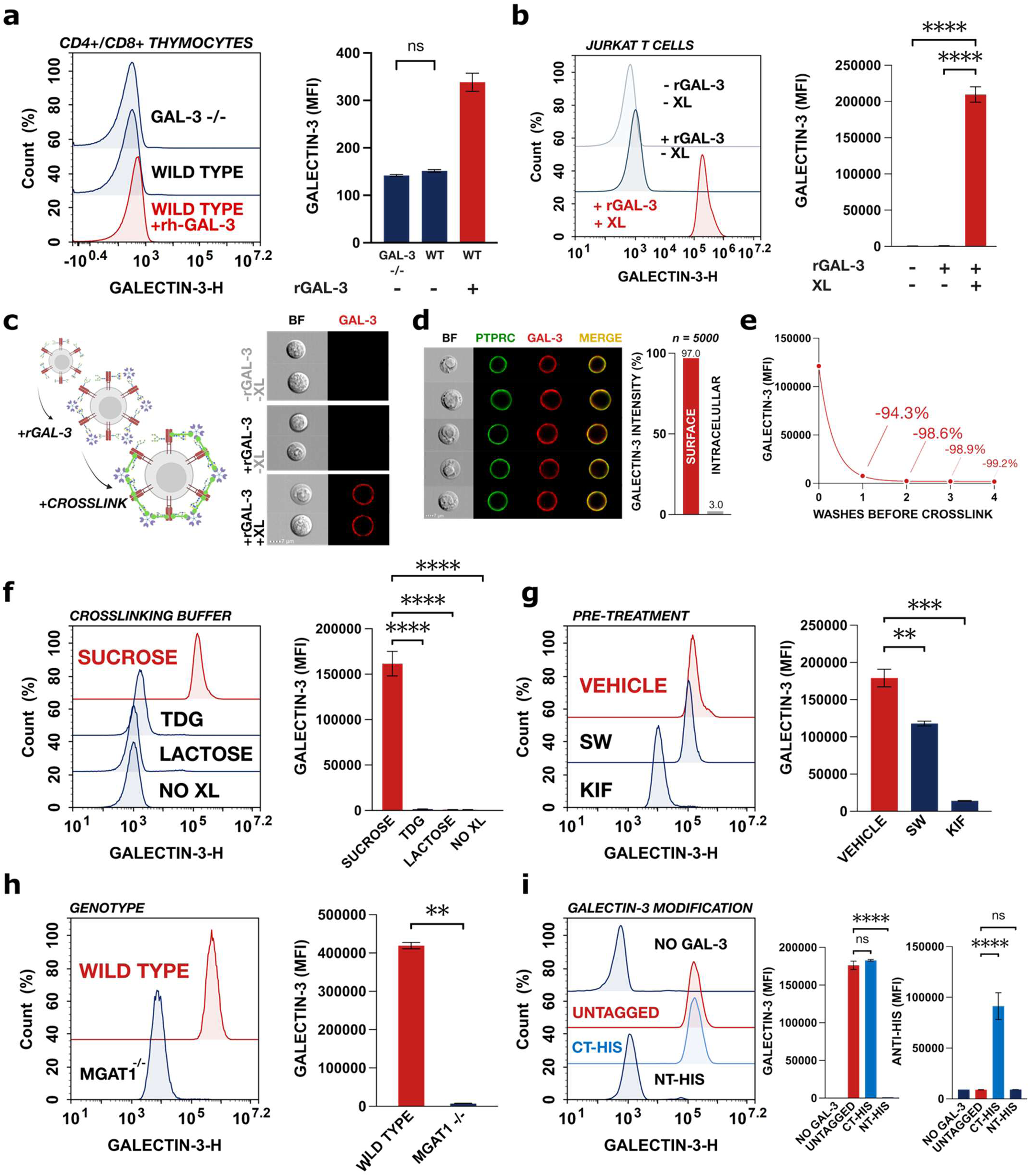
Chemical crosslinking enables robust, specific capture of surface galectin-3. **a** Flow cytometry of galectin-3 staining from *ex vivo* wild type and *Lgals3^-/-^*thymocytes, with and without incubation of recombinant galectin-3 (rGAL-3) as indicated, gated on CD4+/CD8+ DP thymocytes. **b** Flow cytometry of galectin-3 staining on wild type Jurkat cells treated with or without 1.25 µg/ml rGAL-3 and 2 mM DSG as indicated. **c** Workflow schematic for immediate chemical crosslinking of rGAL-3 on Jurkat cells and representative flow cytometry images (n=5000) of Jurkat cells processed as in (b). **d** Representative flow cytometry images of Jurkat cells following crosslinking of rGAL-3 and corresponding surface localization of galectin-3 signal. Galectin-3 (red) surface localization was defined by masking against PTPRC (green). **e** Jurkat cells were crosslinked with 2 mM DSG after 0-4 washes with 200 µL of PBS, galectin-3 staining quantified by flow cytometry. **f** Flow cytometry of immediate crosslinking in the presence of 50 mM sucrose, 50 mM β-lactose or 10 mM thiodigalactoside (TDG). **g** Flow cytometry of galectin-3 crosslinking following treatment of Jurkat cells with 500 nM Swainsonine (SW) or 10 µM Kifunensine (KIF) for 96 hours. **h** Flow cytometry of galectin-3 crosslinking on wild type versus MGAT1 deficient Jurkat cells. **i** Flow cytometry of galectin-3 staining on Jurkat cells after crosslinking of untagged, C-terminal his-tagged (CT-HIS), and N-terminal his-tagged (NT-HIS) galectin-3. Data for **(a-i)**, are representative of ≥ 3 independent experiments. Statistical comparisons by Welch’s t-test **(a, h)** and ordinary one-way ANOVA with Dunnett’s post-hoc test (**b, f-g, i)** and (** = p ≤ 0.01, *** = p ≤ 0.001, **** = p ≤ 0.0001).

To confirm this, we incubated Jurkat T cells with rGAL-3 and performed immediate chemical crosslinking with the homobifunctional, amine-reactive crosslinker disuccinimidyl glutarate (DSG). Crosslinking produced a striking (> 100-fold) increase in surface galectin-3 signal over non-crosslinked controls by conventional flow cytometry and enabled clear detectability by imaging flow cytometry (Fig. 1b-c). Co-staining with CD45 (PTPRC) confirmed that the captured signal was surface-localized (Fig. 1d). Delaying crosslinking until after one, two or four washes with 200 µl PBS reduced signal by >94%, >98% and >99%, respectively (Fig. 1e). Thus, most detectable surface-associated galectin-3 in Jurkat cells is lost during washing unless stabilized immediately.

To distinguish glycan-dependent capture from nonspecific tethering, we performed crosslinking in the presence of soluble galectin ligands. β-Lactose and thiodigalactoside (TDG) inhibited capture, whereas D-sucrose did not (Fig. 1f). Consistent with dependence on physiological N-glycan ligands, DSG capture on Jurkat cells was reduced ∼93% by pretreatment with the α-mannosidase I inhibitor kifunensine (KIF) and ∼33% by the α-mannosidase II inhibitor swainsonine (SW), the smaller SW effect being consistent with compensatory poly-N-acetyllactosamine extension (Fig. 1g, Fig. S1f)^45,46^. As an orthogonal and more complete abrogation of branching, we generated MGAT1-knockout (KO) Jurkat cells by CRISPR editing. MGAT1-KO cells were negative for the branching-dependent lectins L-PHA (*Phaseolus vulgaris leukoagglutinin*, β1,6-branch reactive), E-PHA (*Phaseolus vulgaris erythroagglutinin*, bisecting GlcNAc reactive), and DSL (*Datura stramonium lectin*, β1,4-branch reactive), supporting effective disruption of complex N-glycan biosynthesis (Fig. S1g). By flow cytometry, galectin-3 capture by crosslinking on MGAT1-KO cells was reduced by 97% relative to wild-type Jurkat cells (Fig. 1h). Together, these data establish that (i) crosslinking captures weak but highly specific cell surface galectin-3 interactions, and (ii) galectin-3 engagement of the Jurkat T cell surface is dependent on N-glycan branching.

We next examined whether modification of galectin-3 itself alters capture. Galectin-3 binding has previously been shown to tolerate C-terminal but not N-terminal modification^24^. Indeed, by chemical crosslinking, C-terminally His-tagged galectin-3 was captured nearly identically to untagged protein, while N-terminally His-tagged galectin-3 was not effectively captured at standard doses (Fig. 1i). Intriguingly, titrating N-terminally tagged galectin-3 to higher concentrations resulted in appreciable signal, and this capture remained lactose-inhibitable (Fig. S1h). Finally, we characterized crosslinker concentration, rGAL-3 concentration and reaction time for galectin-3 capture. DSG concentrations of at least 1 mM produced the highest MFIs, although lower concentrations also improved detection relative to non-crosslinked controls (Fig. S1i). rGAL-3 titration produced dose-dependent increases in signal but reduced cell viability at or above 5 µg/ml when added to cells in 96 well plates for flow cytometry experiments (Fig. S1j-k). While longer reaction times increased signal, a 20-min crosslinking reaction balanced capture with preservation of light-scattering characteristics and thus was used subsequently (Fig. S1l).

### Crosslinking enables profiling of global endogenous galectin-3 across immune cells and tissues

We next tested whether immediate crosslinking enables detection of *endogenous* galectin-3 in primary cells directly isolated from lymphoid tissue. Thymocytes were isolated by mechanical dissociation either in standard medium or in medium containing DSG. Resolution of double negative 1 (DN1), DN2, DN3, DN4, DP, CD4⁺ SP, and CD8⁺ SP populations was comparable between preparations, indicating that these populations remain resolvable with the tested marker panel (Fig. 2a, Fig. S2a-c). Immediate crosslinking yielded clear detection of endogenous galectin-3 across all thymocyte populations, unlike non-crosslinked controls (Fig. 2b, Fig. S2d-f). As on Jurkat cells, capture was inhibited by β-lactose but not by D-sucrose (Fig. 2c), and signal was absent in *Lgals3^-/-^* thymocytes (Fig. 2d). Confocal microscopy similarly showed signal in wild type but not *Lgals3^-/-^* thymocytes, with galectin-3 on wild type cells co-localizing with surface Ptprc (Fig. 2e).

**FIGURE 2:**
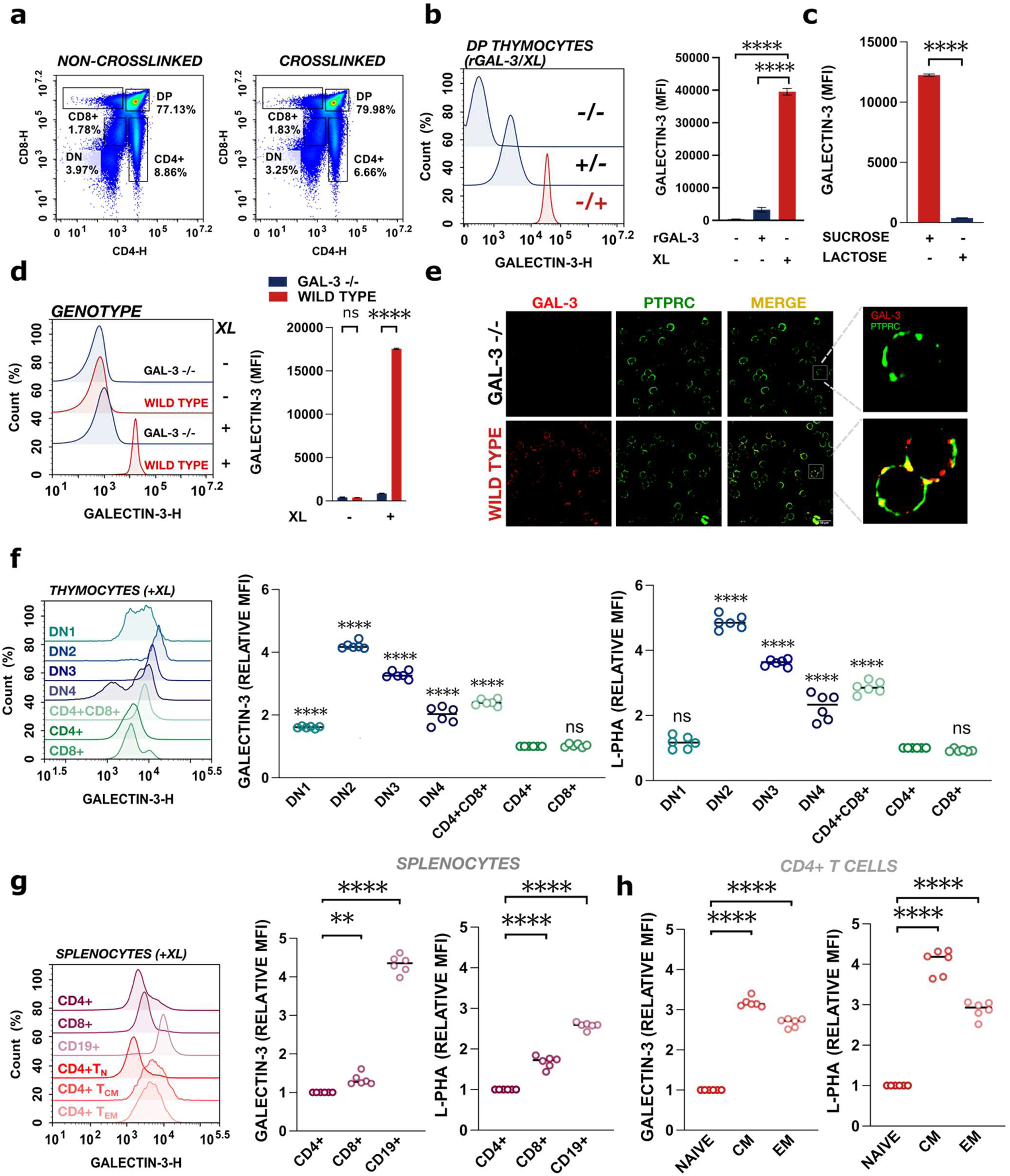
Crosslinking enables profiling of global endogenous galectin-3 across immune cells and tissues. **a** Representative flow cytometry density plots of CD4 vs. CD8 staining on mouse thymocytes before and after crosslinking. **b** Flow cytometry of galectin-3 staining gated on CD4+/CD8+ DP thymocytes with or without 1.25 µg/ml rGAL-3 and 2 mM DSG as indicated. **c** Flow cytometry of crosslinking endogenous galectin-3 in the presence of 50 mM lactose or 50 mM D-sucrose **d** Flow cytometry of galectin-3 signal with or without crosslinking, on *ex vivo* thymocytes from wild type or galectin-3 knockout (*Lgals3^-/-^*) mice, gated on CD4+/CD8+ DP thymocytes. **e** Representative confocal microscopy images of *Lgals3^-/-^* and wild type thymocytes after crosslinking (red = galectin-3, green = Ptprc). **f-g** Flow cytometry of endogenous galectin-3 and L-PHA binding across thymocyte (f) and splenic (g) cell subtypes after crosslinking. Median fluorescence intensity (MFI) values were normalized within each mouse to CD4+ SP values. **h** Flow cytometry of endogenous galectin-3 and L-PHA binding on naive, central memory, and effector memory CD4+ splenic T-cells, normalized within each mouse to naive values. Data for **(a-d)** are representative of ≥ 3 independent experiments. *Ex vivo* immunophenotyping experiments **(f-h)** were performed on young (7 ± 2 weeks) wild type C57BL/6 mice (n = 6; 4 female + 2 male). Statistical comparisons by ordinary one-way ANOVA with Dunnett’s post-hoc test **(b, f-h)** and Welch’s t-test **(c-d)** (** = p ≤ 0.01, **** = p ≤ 0.0001).

We then used immediate crosslinking to survey endogenous galectin-3 across immune cell and tissue types. Because galectin-3 prefers β1,6-branched N-glycans, we predicted that endogenous surface galectin-3 would broadly track with branching as measured by L-PHA. In the thymus, DN2-DP thymocytes carried the highest endogenous galectin-3, while CD4⁺ and CD8⁺ SP thymocytes carried lower but clearly detectable signal (Fig. 2f, Fig. S2b). In spleen and popliteal lymph node (PLN), B cells (CD19⁺) displayed the highest endogenous surface galectin-3, with CD8⁺ T cells slightly exceeding CD4⁺ T cells (Fig. 2g, Fig. S2e-f). Within splenic T cells, CD4⁺ central memory (CD62L^hi^CD44^hi^) and effector memory (CD62L^lo^CD44^hi^) cells carried significantly more endogenous galectin-3 than naïve (CD62L^hi^CD44^lo^) counterparts (Fig. 2h). Among CD8⁺ T cells, effector memory cells were significantly elevated while central memory cells trended upward (Fig. S2g). While endogenous galectin-3 levels were broadly congruent with differences in β1,6-branching detected by L-PHA, we observed that L-PHA was not entirely predictive of endogenous galectin-3. For instance, the relative difference in L-PHA signal was smaller than the difference in galectin-3 signal between splenic B-cells and T-cells. Furthermore, while L-PHA was decreased in CD8⁺ effector memory relative to CD8⁺ naïve T-cells, galectin-3 was elevated. This suggests that although β1,6-branching is a major contributor to galectin-3 lattice organization, additional factors such as galectin-3 availability/distribution may promote or limit galectin-3 binding endogenously. By revealing differences that are otherwise diminished to the point of undetectability without crosslinking, these analyses provide robust and reproducible quantification of endogenous cell surface galectin-3 across thymic and peripheral immune populations.

### xGL-APMS enables enrichment of native surface galectin-3 lattices

We next asked whether chemical capture could be extended into an affinity purification workflow that proteomically identifies surface galectin-3 interactors. Indeed, chemical crosslinking approaches coupled to co-purification and western blotting have previously identified TCR and PTPRC as galectin-3 binding partners^12,43,44^. We thus designed Crosslinking-assisted Galectin Lattice Affinity Purification Mass Spectrometry (xGL-APMS) around five steps: (i) immediate crosslinking of surface galectin-3; (ii) binding and crosslinking of an enrichable anti-GAL-3 IgG; (iii) denaturing lysis to solubilize crosslinked complexes and limit nonspecific binding; (iv) affinity purification; and (v) digestion followed by mass spectrometry. (Fig. 3a).

**FIGURE 3:**
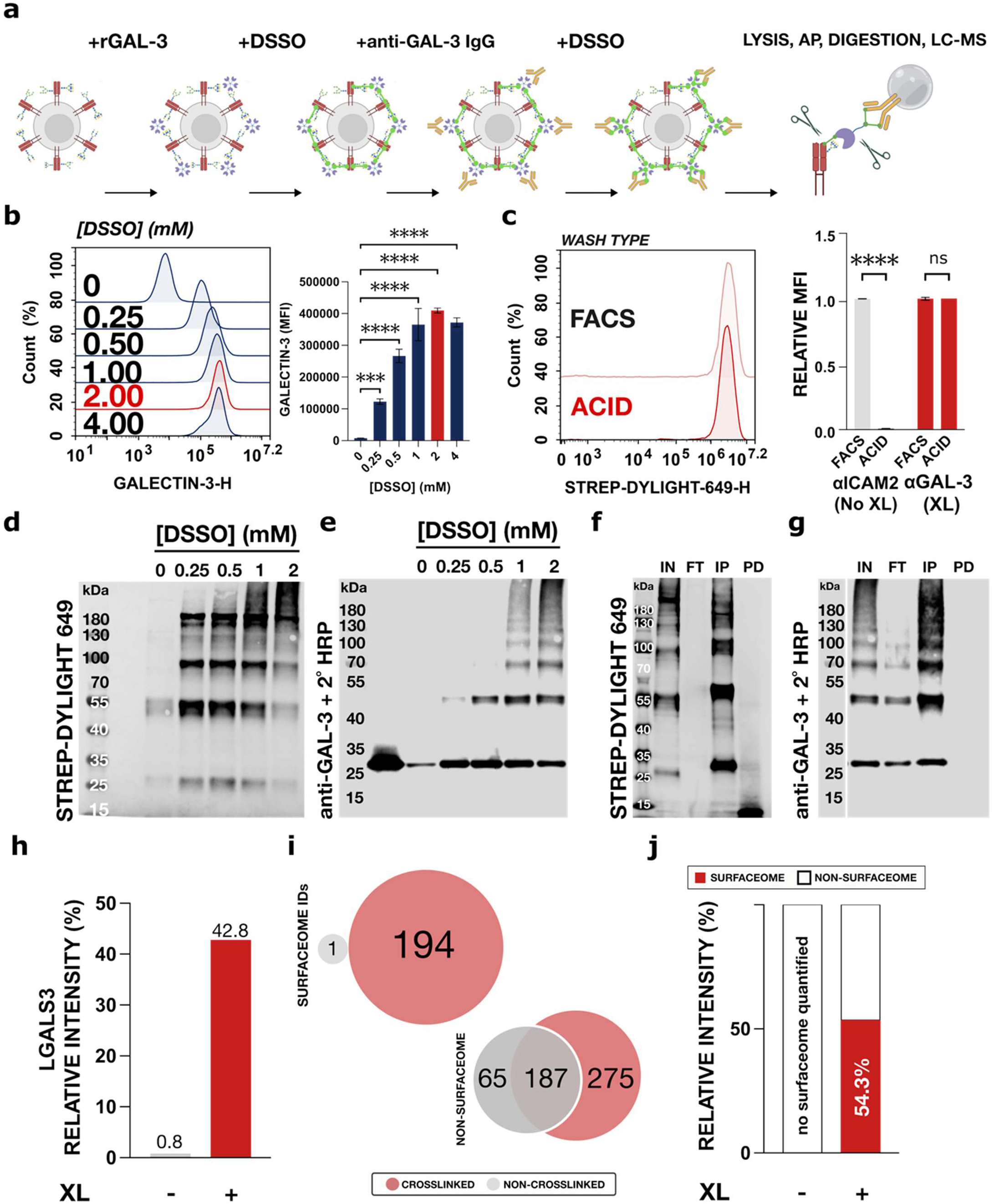
xGL-APMS enables enrichment of native surface galectin-3 lattices. **a** xGL-APMS workflow schematic for crosslinking of recombinant galectin-3 (rGAL-3) and biotinylated anti-GAL-3 on Jurkat cells. Crosslinked cells are lysed in harsh denaturing conditions followed by enrichment on streptavidin agarose, LysC + trypsin digestion, and liquid chromatography mass spectrometry (LC-MS) protein identification/quantification. **b** Flow cytometry of galectin-3 capture on Jurkat cells across a titration of disuccinimidyl sulfoxide (DSSO). **c** Flow cytometry of acid (150 mM NaCl, 20 mM HCl, pH 1.7) washing of surface IgGs anti-ICAM2 (non-crosslinked) and anti-GAL-3 (crosslinked). **d-e** Representative STREP-DYLIGHT-649 **(d)** and anti-GAL-3 IgG + anti-mouse HRP **(e)** immunoblot of whole cell lysate fractions following xGL-APMS preparation across a titration of DSSO for both crosslinking steps. **f-g** Representative STREP-DYLIGHT-649 **(f)** and anti-GAL-3 IgG + anti-mouse HRP **(g)** immunoblot of whole cell lysate (IN), flow-through (FT), purified (IP), and post-digestion (PD) fractions following xGL-APMS preparation. **h** Relative intensity share of galectin-3 (LGALS3) in non-crosslinked vs. crosslinked LC-MS preparations. **i** Venn diagram of surfaceome and non-surfaceome identifications from crosslinked vs. non-crosslinked LC-MS workflows. **j** Relative intensity share of non-surfaceome (white) and surfaceome proteins (red) in non-crosslinked vs. crosslinked LC-MS preparations. Galectin-3 (LGALS3) was excluded from intensity share and count calculations for this analysis. Data shown **(b-g)** are representative of ≥ 3 independent experiments. Statistical comparisons by ordinary one-way ANOVA with Dunnett’s post-hoc test **(b)** and Welch’s t-test **(c)** (*** = p ≤ 0.001, **** = p ≤ 0.0001).

To ensure tractable and economical scaling for proteomic experiments, we examined the effect of increased cell density on galectin-3 binding. At both our previously used dose (1.25 µg/ml) and an upscaled dose (40 µg/ml) of rGAL-3, increasing cell density dramatically reduced galectin-3 MFI, with the flow cytometry scale (1.25 µg/ml rGAL-3, ∼15 million cells/ml) mapping to an upscaled preparation of ∼200 million cells/ml (Fig. S3a-b). We considered whether the observed decrease was due to crosslinker quenching, but further crosslinker titration above 2 mM at high cell density did not improve galectin-3 MFI (Fig. S3c).

For xGL-APMS experiments, we used disuccinimidyl sulfoxide (DSSO), which is chemically similar to DSG but contains an MS-cleavable bond that supports both conventional affinity purification-mass spectrometry and multistage XL-MS^34^. In a direct comparison, DSSO and DSG produced comparable galectin-3 capture by flow cytometry (Fig. S3d). As with DSG, 2 mM DSSO crosslinking produced the highest galectin-3 MFI (Fig. 3b). At 2 mM DSSO, galectin-3 remained surface-bound after β-lactose washing with preserved light-scattering characteristics (Fig. S3e-f). To assess anti-GAL-3 IgG crosslinking efficiency, we performed an acid-stripping assay that removed non-crosslinked anti-ICAM2 IgG but retained crosslinked anti-GAL-3 IgG (Fig. 3c). With increasing DSSO concentrations, high-molecular-weight complexes (> 150 kDa) formed with greater abundance, supporting effective linkage of anti-GAL-3 IgG-galectin-3-glycoprotein complexes by 2 mM DSSO (Fig. 3d, e). Complexes generated under these conditions were effectively enriched using streptavidin-conjugated agarose beads, then digested with trypsin and analyzed by LC-MS (Fig. 3f-g).

Crosslinking increased galectin-3 LC-MS intensity, peptide recovery, and sequence coverage relative to a non-crosslinked workflow (Fig. 3h, Fig. S3h). More strikingly, detection of Jurkat surfaceome proteins was essentially exclusive to crosslinked samples, rising from 1 to 194 surfaceome identifications (Fig. 3i). Excluding galectin-3, surfaceome proteins contributed 54.3% of the summed label-free intensity share in crosslinked samples but no quantifiable intensity in non-crosslinked samples (Fig. 3j). Thus, crosslinking recovers the cell surface galectin-3 lattice, which is otherwise poorly maintained under conventional affinity purification.

### xGL-APMS reveals an N-glycan branching-dependent surface galectin-3 interactome on Jurkat cells

To define galectin-3’s N-glycan branching-dependent surface interactome rigorously, we coupled xGL-APMS to SILAC quantification of wild-type versus MGAT1-knockout (KO) Jurkat cells. Because MGAT1-KO cells lack branched N-glycan ligands and demonstrate dramatically lower galectin-3 binding, they can be mixed 1:1 with WT Jurkat cells and processed in one pot (Fig. S4a-b). This design provides an internally controlled comparison that minimizes technical variance in sample preparation and discriminates glycan-dependent interactors from non-specific contaminants and workflow artifacts.

Two SILAC experiments, each comprising two biological replicates, identified 1,036 proteins across four runs, including 212 surfaceome proteins. Restricting high-confidence hits to those present in ≥ 2 replicates, and enriched ≥ 5-fold in wild type over MGAT1-KO with p ≤ 0.05, yielded 176 proteins, of which surfaceome proteins derived from the SURFY database comprised 125 by count and 96.4% of intensity share^47,48^ (Fig. 4a-d). Alternative surfaceome annotation or filtering for N-glycosylated proteins displayed comparable enrichment^49,50^ (Fig. S4c). While most intracellular proteins did not meet the WT:MGAT1-KO enrichment criteria, some enriched proteins represent membrane proximal components, such as LCK, or plasma membrane proteins that are not captured by surface annotation, such as CD3E, indicating that our analysis likely underestimates the true degree of surfaceome enrichment^51^. The enriched galectin-3 surface client list expands upon Jurkat-expressed galectin-3-associated proteins catalogued in BioGRID 5.0, IntAct and STRING-DB^52–55^ (Fig. 4e).

**FIGURE 4:**
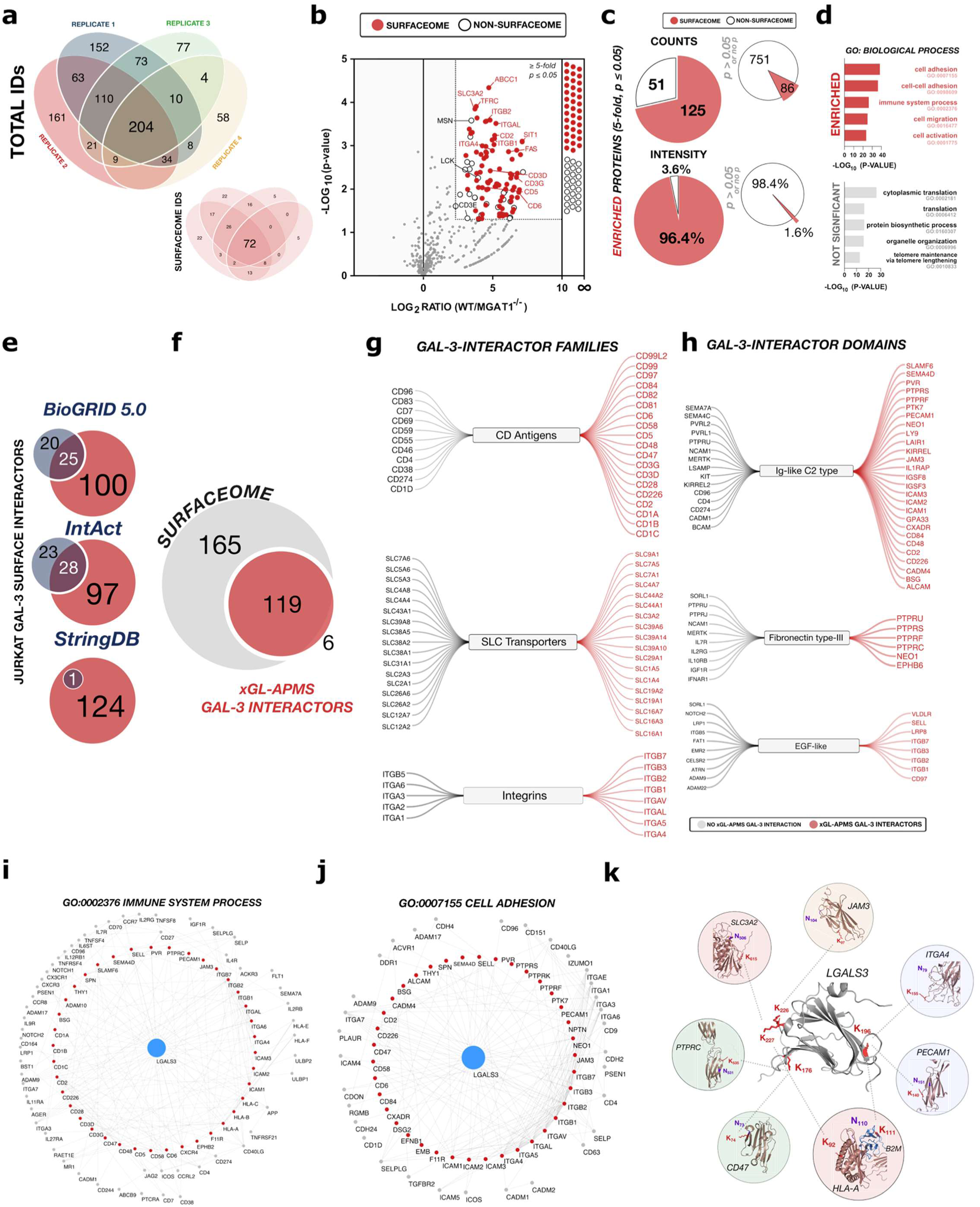
xGL-APMS reveals an N-glycan branching-dependent surface galectin-3 interactome on Jurkat cells. **a** Venn diagram of total protein IDs and surfaceome IDs from four independent SILAC xGL-APMS preparations. Replicates 1 (blue), 2 (red), and 4 (yellow) were performed by labeling wild type Jurkat cells with light arginine and lysine and MGAT1^-/-^ cells with heavy arginine (Arg 10) and lysine (Lys 8). Replicate 3 (green) was performed with reversed SILAC labeling. **b** Volcano plot of protein enrichment (Log₂ wild type vs. MGAT1^-/-^ ratio) in SILAC xGL-APMS experiments. **c** Distribution of surfaceome (red) versus non-surfaceome (white) proteins among significantly enriched hits (5-fold, p ≤ 0.05) and non-significant/non-reproducible proteins (p > 0.05) by count and intensity share. **d** Gene ontology (GO) of top five biological processes (BPs) in the significantly enriched vs. non-significant pools. **e** Venn diagram of Jurkat galectin-3 surface interactors reported in BioGRID 5.0, IntAct, and String DB repositories versus xGL-APMS-identified interactors. **f** Venn diagram of Jurkat surfaceome proteins reported in Sánchez et al., 2026 versus xGL-APMS significant surfaceome hits. **g-h** Classification of xGL-APMS detected galectin interactors and non-interacting surface glycoproteins by protein family **(g)** and repeat domain content **(h). i-j** Physical subnetwork map of xGL-APMS-identified galectin-3 interactors (red) and Jurkat-expressed (TPM ≥ 1) first neighbors under the GO:0002376 Immune System Process **(i)** and GO:0007155 Cell Adhesion **(j)** BP annotations. Non-galectin-3 edges were sourced from String-db with confidence ≥ 0.5. **k** Multistage crosslinking mass spectrometry (MS^2^-MS^3^) interpeptide linkage identifications with corresponding nearest N-glycosylation site mapping by proximity.

To contextualize xGL-APMS-identified galectin-3 interactors, we compared enriched clients to our previously characterized Jurkat surfaceome, obtained by amine-reactive cell surface biotinylation followed by the same denaturing lysis and streptavidin enrichment procedures^48^. Of 284 identified surfaceome proteins identified by cell surface biotinylation (surfaceomics), 119 appeared among xGL-APMS galectin-3 surface interactors while 165 did not (Fig. 4f), indicating that galectin-3 engages a large but bounded subset of the surfaceome rather than the entire surface indiscriminately. Functional family and repeat domain annotation reinforced this selectivity. CD antigens, SLC transporters, integrins, tetraspanins, intercellular adhesion molecules (ICAMs), and immunoglobulin superfamily members (IgSFs) were prominent among interactors, whereas interferons, ADAM proteases, and semaphorins were depleted or absent (Fig. 4g, Fig. S4d). By repeat domain, immunoglobulin (Ig)-like, Fibronectin-type III, EGF-like, and tyrosine phosphatase domains were abundant while Mannose-6-phosphate receptor homolog (MRH), glycine-serine rich (GS), and complement subcomponent (CUB) domains were absent (Fig. 4h, Fig. S4d). We visualized the galectin-3 surface interactome in context by mapping interactors and their first neighbors onto STRING-DB networks for GO:0002376 (immune system process) and GO:0007155 (cell adhesion) (Fig. 4i-j).

We also asked whether N-glycosylation site number distinguished interactors from the broader surfaceome. Comparing xGL-APMS interactors to non-interactors, a weak preference for medium N-glycan number receptors was observed, but interactors spanned the full N-glycan number range, including receptors with few N-glycan sites (Fig. S4e). We interpret this to mean that in a high-branching cell such as Jurkat^20^, where galectin-3-preferred branched glycoforms are abundant, a single/few well-presented sites can suffice. These observations suggest that glycoform identity and spatial presentation may contribute to engagement beyond potential N-glycosylation-site count. N-glycan number may become a stronger determinant of galectin-3 binding as branching decreases, for example in low-branching cell states or in *MGAT5* deficiency.

### Site-specific interlinks support direct interaction and define the sensitivity limits of XL-MS

The xGL-APMS based workflow inherently permits multistage crosslinking mass spectrometry (MS^n^/XL-MS) to site-specifically probe galectin-3-glycoprotein interactions. Such data provide additional confidence for direct interaction and yield structural insight into individual galectin-3-ligand interfaces. MS^2^-MS^3^ acquisition on xGL-APMS samples identified nine interlinks involving galectin-3, establishing feasibility (Fig. 4k). As expected, every interlinked partner was also identified at the protein level by xGL-APMS. We identified interlinks involving one lysine site (K176) near the carbohydrate binding pocket of galectin-3, and two additional binding pocket-adjacent crosslinked sites (K196, and K226/227) in similar frequency (Fig. S4f-g). Four additional lysines are present in the CRD but were not identified in interlinks^56^.

Notably, interlinks were recovered for only a subset of xGL-APMS interactors, raising two competing explanations: i) that few xGL-APMS-identified interactors are direct, or ii) that XL-MS is sensitivity-limited. In the case of the latter, we hypothesized that interlink identification should be biased toward abundant interactors. Indeed, this was the case by both transcript abundance (TPM) and MS intensity (Fig. S4h). We then tested this directly by asking whether an interactor identified by xGL-APMS but *not* by interlink is nonetheless a direct, glycan-dependent binder. Indirect or bystander capture should persist regardless of glycosylation status, whereas a direct glycan-dependent interaction should be abolished by NXS/T mutation. We selected CD5 for its high xGL-APMS intensity and tractable mutability (two N-glycosylation sites), generating wild-type (HA-CD5^WT^) and N-glycan-deficient (HA-CD5^N116A,N241A^) constructs. After confirming comparable expression, surface localization, and the expected glycosylation states (Fig. S4i-l), immunoblotting showed that co-purification of galectin-3 and CD5 was dependent on its N-glycosylation (Fig. S4m). CD5 is therefore a glycan-dependent galectin-3 ligand despite the lack of interlink identification, supporting that XL-MS sensitivity likely underreports galectin-3 interactors.

### Orthogonal and multi-faceted validation of the xGL-APMS galectin-3 interactome

To validate xGL-APMS interactions more broadly, we probed galectin-3 surface interactions using a series of orthogonal approaches. We first examined co-purification by immunoblotting, using the thiol-cleavable amine-reactive crosslinker DSP, which captured galectin-3 comparably to DSG and DSSO and allowed recovery of monomer bands after reduction (Fig. S4k). PTPRC and TFRC were detected in the purified fraction, whereas β-actin was not detected (Fig. 5a). Both receptors also remained in the flow-through, consistent with intracellular pools, incomplete capture, or enrichment of only part of the surface pool (Fig. 5a).

**FIGURE 5:**
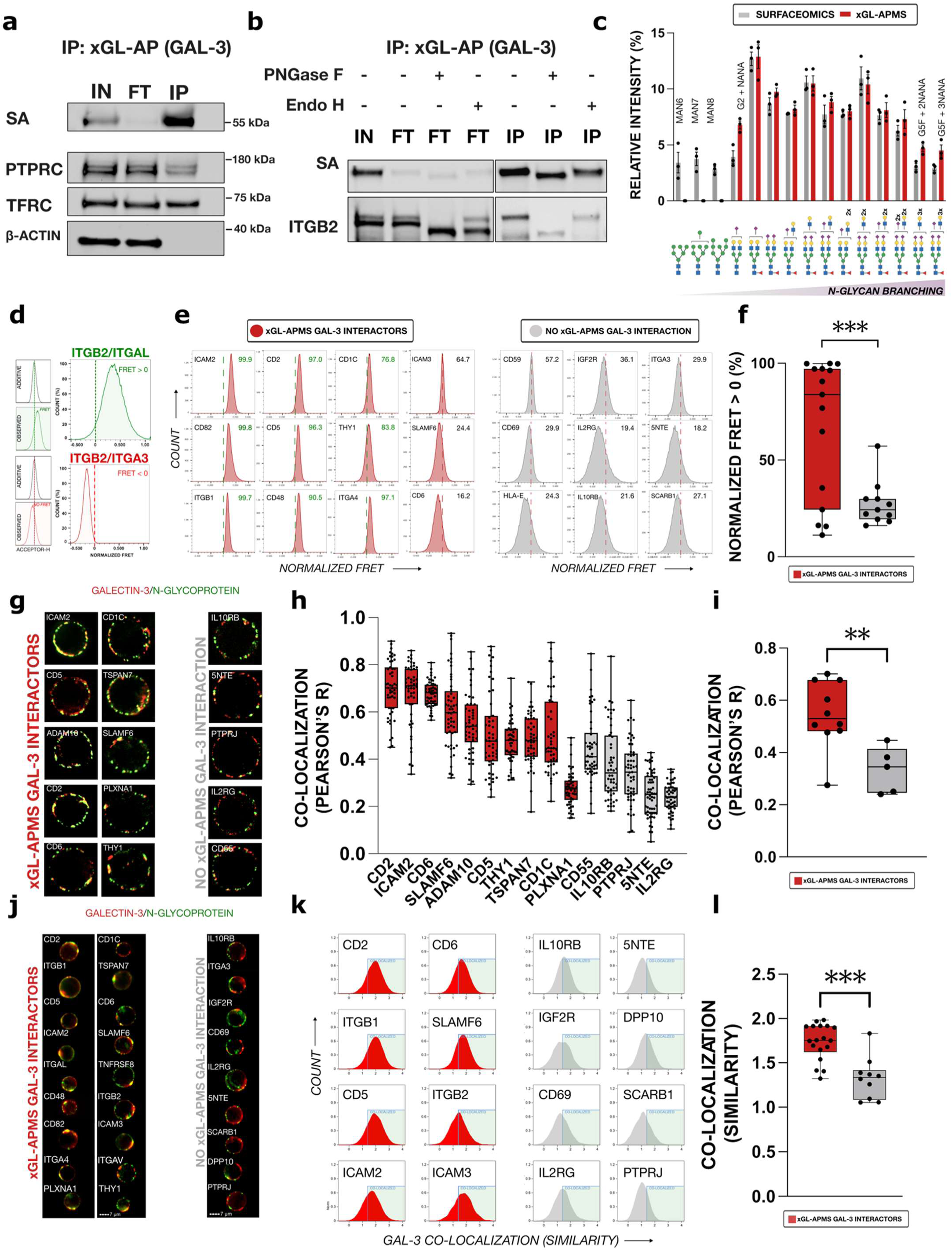
Multi-faceted validation of the xGL-APMS galectin-3 interactome. **a** Representative DSP immunoblot of whole cell lysate (IN), flow-through (FT), and purified (IP) fractions from wild type Jurkat cells under the xGL-APMS workflow, probed by STREP-DYLIGHT-649 (SA), anti-PTPRC, anti-TFRC, and anti-β-Actin. **b** Representative immunoblot of ITGB2 prepared under the xGL-APMS workflow with DSP crosslinking and corresponding assessment of glycosylation by PNGase F and Endo H treatment. **c** MALDI-ToF glycomic comparison of relative N-glycan intensities under untargeted surfaceomics (grey) and xGL-APMS (red). Relative abundance was calculated as the ratio of individual glycan peak height to total peak sum across 15 reproducibly identified N-glycoforms (n=3). Non-overlapping enriched and depleted N-glycoforms are labeled with shorthand nomenclature. **d** Flow-FRET calculation for a tightly interacting surface heterodimer (ITGB2-ITGAL) and non-interacting surface heterodimer (ITGB2-ITGA3). **e-f** Flow-FRET analysis of galectin-3 and 26 Jurkat surface proteins (15 xGL-APMS galectin-3 interactors and 11 lacking xGL-APMS interaction). **g** Representative confocal microscopy images (n = 50 cells) of galectin-3 (red) and 15 Jurkat surface proteins (green, 10 xGL-APMS interactors and 5 lacking xGL-APMS interaction). **h-i** Galectin-3 co-localization by confocal microscopy for analyzed Jurkat surface proteins **(h)** and comparison of xGL-APMS galectin-3 interactors and non-interactors **(i). j** Representative flow cytometry capping images (n = 2500) of galectin-3 (red) and Jurkat surface receptors (green). **k-l** Galectin-3 co-localization (Similarity) histograms of select xGL-APMS interactors. Co-localization minimum (blue) of 1.4 indicated **(k)** and corresponding comparison of xGL-APMS galectin-3 interactors (red) and non-interactors (grey). Statistical comparisons performed by Welch’s t-test (** = p ≤ 0.01, *** = p ≤ 0.001).

We distinguished these possibilities for ITGB2, an xGL-APMS galectin-3 interactor whose surface and intracellular pools in Jurkat cells are separable by Endo H sensitivity^48,57,58^. The xGL-APMS-purified ITGB2 fraction was Endo H resistant and PNGase F sensitive, indicating mature N-glycans consistent with a post-Golgi surface pool. The flow-through, while dominated by the Endo H-sensitive (immature, intracellular) species, also retained the Endo H-resistant high molecular weight band at levels comparable to input (Fig. 5b). Galectin-3 therefore likely engages only a subset of the *surface* pool of its ligands. Consistent with this, titrating galectin-3 induces cell death well before surface binding saturates, indicating that not all available surface ligands are engaged at one time (Fig. S1j).

We next compared glycans recovered by xGL-APMS with those obtained by untargeted surface enrichment (surfaceomics). MALDI-TOF glycomics identified three complex N-glycoforms reproducibly enriched by xGL-APMS across three replicates (G2+NANA, G5F+2NANA, G5F+3NANA), plus five additional branched N-glycoforms detected exclusively in xGL-APMS preparations in one or two replicates (Fig. 5c, Fig. S5a-b). Conversely, high-mannose N-glycoforms were not detected in xGL-APMS preparations but were detectable in all three surfaceomic samples. xGL-APMS thus reproducibly captures a glycomically distinct, branching-enriched subset of the surfaceome.

Individual interactors and non-interacting surface proteins were then validated using Förster resonance energy transfer (FRET), confocal co-localization, and antibody-induced capping-mediated redistribution. FRET was measured using a PE^DONOR^-APC^ACCEPTOR^ antibody-based flow-FRET approach^59–61^. Here, we compared the additive spectral bleed into the acceptor (667/30 nm) channel from PE and APC single-stained controls versus observed acceptor signal when co-staining with PE and APC-conjugated antibodies. For a tightly interacting pair such as LFA-1 (ITGB2-ITGAL), positive FRET was detectable by rightward shifts in the co-stained acceptor channel relative to additive spectral bleed of single-stained controls, resulting in positive normalized FRET. By contrast, a non-interacting integrin pair (ITGB2-ITGA3) served as a negative control, producing no detectable normalized FRET, as expected (Fig. 5d)^62–64^. Because FRET is spatially stringent, true interactions can be missed when antibody epitopes are too distant. We therefore tested a broad panel of 26 surface proteins (15 xGL-APMS interactors, 11 non-interactors) as FRET donors to galectin-3 (Fig. S5c). Definitively positive flow-FRET was detected for 9 of 15 xGL-APMS interactors, with an additional interactor (ICAM3) displaying low but positive signal. Conversely, none of the 11 non-interactors tested displayed a definitive FRET signal (Fig. 5e). As a group, interactors displayed significantly higher FRET frequency than non-interactors (Fig. 5f).

By high-resolution confocal microscopy of 15 surface proteins (10 interactors, 5 non-interactors), galectin-3 interactors showed significantly greater co-localization with galectin-3 than non-interactors across 1 µm sections (Pearson’s R) (Fig. 5g-i, Fig. S5d). Finally, we exploited galectin-3 capping (the redistribution of surface galectin-3 into discrete patches upon anti-GAL-3 IgG addition at 37 °C for 30 min) followed by crosslinking, immunofluorescent staining, and imaging flow cytometry, to ask whether interactors are drawn into patches with galectin-3^65–67^ (Fig. S5e-f). Across 28 surface proteins, interactors showed significantly greater co-mobility mediated co-localization with galectin-3 than non-interactors (Fig. 5j-l, Fig. S5g-h).

Across the validation panel we examined a total of 35 surface proteins spanning the breadth of surfaceome abundance (24 xGL-APMS interactors and 11 non-interactors) using six complementary approaches: N-glycosite mutation, immunoblotting, MALDI-ToF glycomics, flow-FRET, confocal co-localization, and imaging flow cytometry of capped cells. The -omics and immunoblotting data established that xGL-APMS reproducibly enriches a defined, glycomically distinct subset of the surface proteome, while the non-omics approaches independently corroborated individual interactions and support the stringency of our interactor definition criteria.

### xGL-APMS unveils the endogenous galectin-3 interactome on primary mouse thymocytes

Galectin-3 interactomes reported to date have relied on exogenous galectin-3, which may differ from endogenous galectin-3 in concentration, tissue/microenvironment distribution, and/or protein isoform. Because xGL-APMS requires no exogenous, tagged, or overexpressed bait, we reasoned it could enable proteomic interrogation of endogenous galectin-3 interactions. Before pursuing this, we sought to use crosslinking-enabled galectin-3 detection to compare endogenous galectin-3 to exogenously added rGAL-3 both quantitatively and in its organization at the cell surface.

We first examined the surface distribution of exogenous galectin-3 across a titration of rGAL-3 concentrations on Jurkat cells by imaging flow cytometry. At high doses, galectin-3 staining was ring-like, spanning most or all of the cell circumference. Interestingly, at doses below 1.25 µg/ml, staining appeared to resolve into discrete, high-intensity puncta separated by regions of distinctly dimmer signal. Quantification of perimeter coverage and uniformity confirmed this impression, with the perimeter length bearing detectable galectin-3 signal and staining uniformity increasing progressively with dose (Fig. 6a). These observations are consistent with preferential association in discrete regions at lower concentrations and broader surface coverage as galectin-3 dose increases.

**FIGURE 6:**
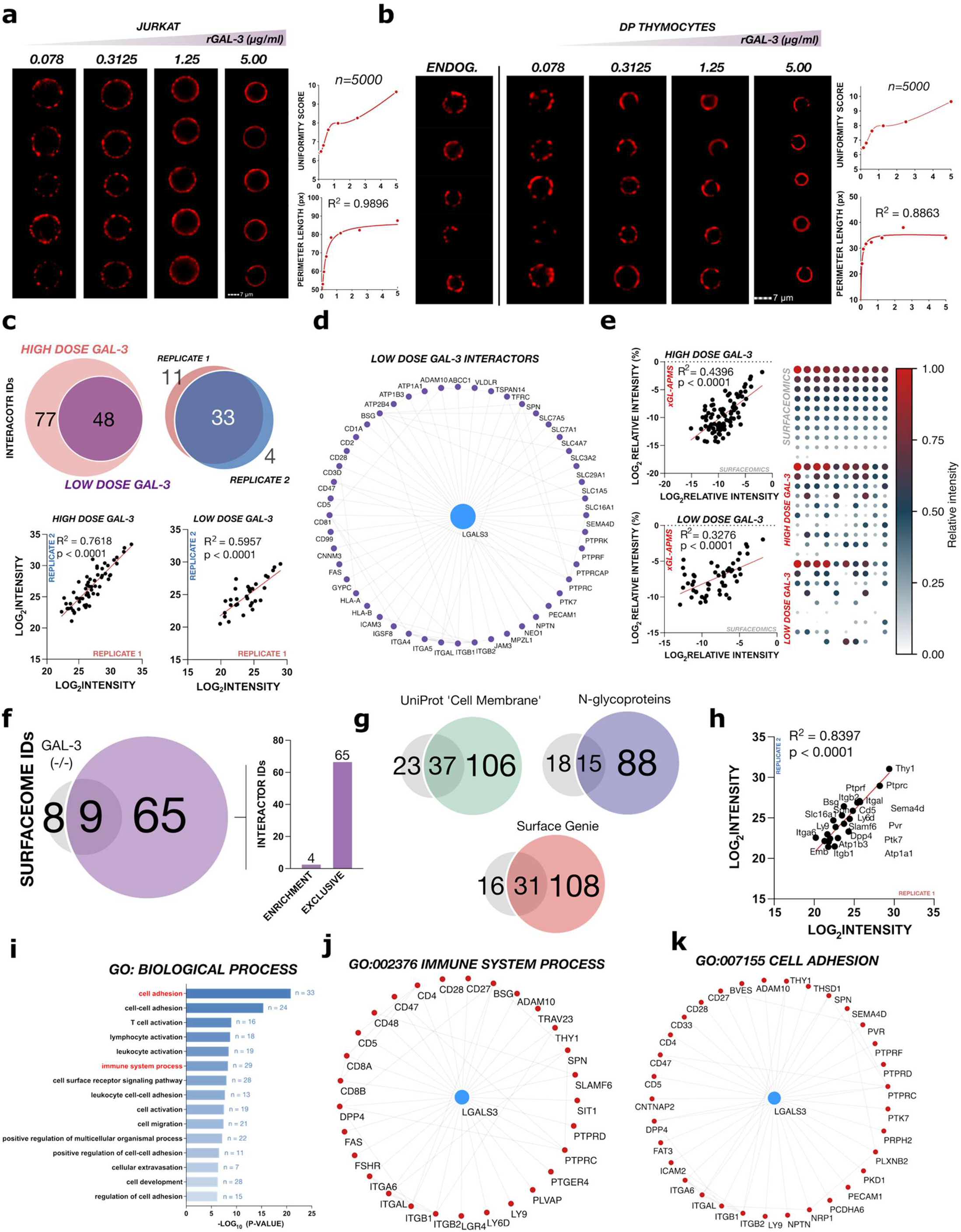
xGL-APMS reveals the endogenous galectin-3 interactome on primary mouse thymocytes. **a** Representative flow cytometry images (n = 5000) of Jurkat cells following a titration of rGAL-3 and corresponding quantification of uniformity (via the *Homogeneity* feature) and perimeter length. **b** Representative flow cytometry images (n = 5000) of primary mouse thymocytes cells following immediate crosslinking for capture of endogenous galectin-3 or a titration of rGAL-3 and corresponding quantification of uniformity and perimeter length**. c** Venn diagram of Jurkat galectin-3 interactors identified by previous SILAC xGL-APMS experiments (high dose) vs. step-down low dose galectin-3 SILAC experiments and linear regression of interactor intensities quantified under high dose and low-dose galectin-3 experiments. **d** Physical subnetwork map of Jurkat galectin-3 interactors identified in step-down low dose galectin-3 SILAC experiments. **e** Linear regression analysis and intensity-arranged array of xGL-APMS interactor quantification with high dose galectin-3 and low dose galectin-3 vs. untargeted cell surface proteomics (surfaceomics). **f** Venn diagram of SURFY-derived surfaceome proteins identified from wild type vs. galectin-3 knockout (Lgals3^-/-^) thymocytes under xGL-APMS preparations. **g** Wild type vs (Lgals3^-/-^) identification venn diagrams filtered by three additional databases (Uniprot ‘Cell Membrane’ annotation, Uniprot N-glycosylation, and Surface Genie with SPC score ≥ 1). **h** Linear regression analysis of xGL-APMS interactor intensities from independent biological replicates. **i** Top GO: Biological Process terms enriched in the endogenous galectin-3 interactome. **j-k** Interactome maps of highlighted GO: Biological Process terms in the endogenous galectin-3 interactome. Non-galectin-3 edges were sourced from StringDB physical subnetwork data with confidence threshold ≥ 0.5.

Endogenous galectin-3 on primary DP thymocytes also showed discrete and punctate staining, like the low-dose rGAL-3 pattern on Jurkat cells. Addition of increasing doses of exogenous galectin-3 to thymocytes progressively converted this pattern to the ring-like distribution observed on Jurkat cells with higher rGAL-3 doses (Fig. 6b). Thus, supraphysiologic galectin-3 qualitatively alters how galectin-3 is arranged at the cell surface.

These findings highlighted the importance of extending xGL-APMS to a purely endogenous galectin-3 interactome. However, the potentially large quantitative difference between endogenous and supraphysiological MFIs raised concerns for proteomic sensitivity. We therefore explored the sensitivity limit of the xGL-APMS approach with a step-down SILAC experiment in Jurkat cells, in which cells at high density (200 million cells/ml) incubated with high-dose (40 µg/ml) or low-dose (1.25 µg/ml) rGAL-3 were crosslinked, then mixed 1:1 prior to subsequent xGL-APMS processing (Fig. S6a). This design enabled direct comparison of interactor intensities across doses and facilitated precursor detection through the paired high-dose sample^68^. Under this framework, 48 of the original 125 galectin-3 interactors were quantified at low dose, with strong qualitative and quantitative reproducibility (Fig. 6c-d, Fig. S6b-c).

This experiment additionally revealed a dose-dependent shift in interactor selection. At high dose, galectin-3 interactor intensities correlated moderately with surfaceomic intensities, indicating that surface abundance partially drives apparent galectin-3 interaction. At low dose, this correlation was weakened (Fig. 6e), suggesting that as galectin-3 approaches physiological levels, surface abundance becomes a less important determinant of lattice incorporation. This difference further establishes the importance of a purely endogenous approach.

We thus proceeded to capture the endogenous native cell surface galectin-3 interactome on primary mouse thymocytes. To do so, we crosslinked endogenous galectin-3 directly in freshly harvested primary tissue, then linked biotinylated anti-GAL-3 IgG, purified crosslinked complexes on streptavidin, digested, and identified interactors by LC-MS. We confirmed by acid stripping that both crosslinked endogenous galectin-3 and crosslinked anti-GAL-3 IgG were retained at the cell surface, indicating effective capture at both steps (Fig. S6d). Because Mgat1 deletion is embryonically lethal and thymocyte-specific deletion of Mgat1 leads to thymocyte death^41,69–71^, we used Lgals3*^-/-^* thymocytes as the negative control throughout. Wild-type and Lgals3*^-/-^* thymocytes displayed comparable CD4⁺ SP, CD8⁺ SP, and DP composition (Fig. S6e), and Lgals3*^-/-^*thymocytes processed through the xGL-APMS workflow yielded low streptavidin signal relative to wild type (Fig. S6f).

Proteomically, galectin-3 was not identified in Lgals3*^-/-^*thymocytes but was abundant in both wild-type replicates (Fig. S6g). Applying enrichment-based filtering (SURFY-derived surfaceome proteins enriched ≥ 5-fold in wild type over Lgals3*^-/-^*), we identified 69 endogenous interactors across two biological replicates. Sixty-five were exclusive to wild-type cells, while four (Atp1a1, Ptprc, Slc16a1, Thy1) were detected in both genotypes but strongly enriched in wild type and thus retained as hits. (Fig. 6f, Fig. S6h-i). Filtering by alternative surfaceome databases (Uniprot ‘Cell Membrane’ annotation, N-glycoproteins, and Surface Genie annotations) demonstrated similar favorability towards wild-type samples^72^ (Fig. 6g). Interactors identified in both replicates were reproducibly quantified by intensity (Fig. 6h). Functional annotation was largely concordant with our Jurkat results. Enriched biological processes and molecular functions centered on cell adhesion, immune system process, cell migration, transport, and cell development, and enriched KEGG and Reactome pathways highlighted cell adhesion molecule (CAM) and IgSF-CAM signaling, T cell receptor signaling and primary immunodeficiency, hematopoietic cell lineage, basigin and integrin interactions, and immunoregulatory interactions (Fig. 6i-k, Fig. S6j-l). These results establish the feasibility of identifying galectin-3 interacting proteins at proteome scale in primary tissues without perturbing the endogenous galectin-3 lattice, and support endogenous galectin-3’s role in T cell adhesion, activation, and development.

## Discussion

Despite galectin-3’s central role in regulating T cell biology, native and endogenous studies of the cell surface galectin-3 lattice have been limited by its inherently labile interactions. Existing approaches to probe native galectin-3 interactions provide complementary information but differ in bait requirements, spatial specificity and sensitivity. Here, immediate chemical crosslinking stabilizes wash-sensitive surface galectin-3 and enables affinity purification of its associated complexes, including those formed by unmodified endogenous galectin-3 on primary cells.

Crosslinking increased global surface galectin-3 signal > 100-fold and raised surfaceome recovery from a single protein to nearly 200. This gain in sensitivity enabled comparison of endogenous galectin-3 glycoprotein association across immune populations and identification of a complex N-glycan-dependent network in Jurkat cells. Complementary biochemical and imaging assays supported selected interactions, and the workflow extended to endogenous galectin-3 in primary thymocytes. In this respect, xGL-APMS is analogous to ChIP-seq for the glycocalyx. Just as formaldehyde crosslinking converted ephemeral transcription factor occupancy into stable, immunoprecipitable signal, immediate chemical crosslinking converts low-affinity, galectin networks into covalently stabilized complexes that survive stringent denaturing lysis, affinity purification, and mass spectrometric identification^31,32^.

Of particular interest is the scale of the galectin-3 interactomes uncovered by xGL-APMS. As a secreted protein that binds common glycoepitopes, we conceptualized that galectin-3 carries the potential for a widespread interactome with influence across a broad range of biological functions. Diverse interactomes are not a universal feature of all lectins, however. Some lectins (e.g., siglecs/selectins) have been found to interact with a relatively restricted set of glycoprotein ligands when examined in physiological contexts^73–75^. Thus, supraphysiological and permissive interactome mapping by existing approaches raised concerns regarding the extent to which the galectin-3 interactome was overestimated. Critically, even under stringent criteria, the galectin-3 interactomes uncovered by xGL-APMS on the Jurkat T cell (125 interactors) and endogenous thymocyte (69 interactors) surface were extensive, supporting the breadth of biological functions which galectin-3 may regulate. However, across xGL-APMS and orthogonal methods, our data did not support universal galectin-3 interaction with all surface N-glycoproteins. This suggests that while widespread, the physiological galectin-3 interactome retains selectivity for specific receptors.

The extensive interactomes recovered by xGL-APMS point to a variety of underexplored binding partners for galectin-3. While previous studies supported galectin-3’s role in regulating T cell adhesion, interactions under the adhesion umbrella were largely limited to integrin complexes (e.g., ITGB1/ITGAV) and cell adhesion molecules (CAMs)^76–79^. In addition to identifying these interactors, xGL-APMS also returned a slate of important cell adhesion complexes, including CD2-CD58 (LFA-3)^80,81^ and PLXNB1-SEMA4D^82^, whose oligomerization or surface retention may intriguingly be regulated by engagement with the galectin lattice. Solute transporters also constituted a major component of the galectin-3 interactome. In addition to the highly abundant amino acid transporter SLC3A2, which has previously been shown to interact with galectin-3^83^, xGL-APMS also identified key transporters responsible for cellular pH regulation via Na^+^/H^+^ ion transport^84^ (e.g., SLC9A1), carbon metabolite transport^85^ (e.g., SLC16A1), and the central zinc ion transport complex responsible for driving epithelial to mesenchymal transitions^86,87^ (SLC39A6-SLC39A10). Lastly, while galectin-3’s role in regulating immune function has previously centered around its engagement with the T-cell receptor (TCR) complex and a small subset of auxiliary immunomodulators^12,43,88^ (e.g., PTPRC, LAG3), xGL-APMS data supported that galectin-3 engages additional immunomodulatory proteins, including both co-stimulatory^89–91^ (e.g., SLAMF6, CD82, CD48) and co-inhibitory^92–94^ (e.g., CD5, SIT1, LAIR1) receptors. Thus, galectin-3’s regulation of T-cell activation may integrate many different proteins rather than exclusively regulating TCR and its most well characterized immunomodulators. Interrogation of galectin-3’s role in regulating surface localization and retention of these interactors would expand the landscape of galectin-3 biology and is a focus of ongoing efforts.

A key limitation of xGL-APMS in its current form is its inability to definitively distinguish direct from indirect interactions, a challenge shared by other interaction mapping approaches. Highlighting this limitation, we identified interactors such as beta-2 microglobulin (B2M) and CD81 that are not known to carry N-glycans, potentially reflecting capture via glycan-independent contact, or indirect association^95,96^. Importantly, in this context indirect association is not necessarily uninformative or equivalent to non-specific capture. The B2M interlink is interpretable via its heterodimer formation with class I MHCs which present a direct N-glycan ligand for galectin-3. Thus, crosslinking may not resolve which subunit of a heterodimer bears the glycan being engaged in all cases, but it does identify that the intact, functional complex is bound by galectin-3. Rather than false positives, such identifications indicate the composition of the functional protein complexes engaged by the galectin-3 lattice.

Targeted glycosylation-site mutation and XL-MS can test whether capture depends on a candidate glycan. Mutation of both CD5 N-glycosylation sites abolished galectin-3 enrichment despite preserved surface expression, supporting N-glycan dependence of xGL-APMS capture. However, this does not by itself exclude indirect effects on conformation or surface organization. XL-MS provides complementary residue-level information but recovered relatively few interlinks in the present implementation. The abundance bias among crosslinked partners is consistent with limited detection sensitivity. Enrichable crosslinkers, extensive fractionation, and additional enrichment steps offer potential routes to deeper interlink coverage. Achieving this depth would be of substantial value because interlinks impose distance constraints on specific lysines. Since lysines near the galectin-3 interface may report on the N-glycan sites mediating contact, XL-MS could nominate the individual glycosites responsible for galectin-3 engagement on multiply glycosylated receptors, thereby identifying potentially targetable galectin-3 interaction sites.

As an approach that does not depend on modified bait, xGL-APMS may be extended to other galectins and glycan-binding proteins for which suitable antibodies are available. Such adaptation could support comparison of associated networks between the galectins. Furthermore, xGL-APMS is inherently compatible with established quantitative frameworks including SILAC, label-free quantification, and isobaric labeling, enabling comparison of lattice composition across biological states. These capabilities broaden the potential application of this workflow in glycobiology. Of translational interest, pairing flow cytometry quantification of surface galectin-3 with xGL-APMS interactomic profiling in patient-derived tissue samples may reveal new candidate biomarkers spanning autoimmunity, fibrosis, heart disease, and cancer.

The composition of T cell galectin-3 lattices had previously been examined in low-throughput or non-physiological contexts. By chemically stabilizing the galectin-3 lattice at the cell-surface before enrichment and LC-MS analysis, xGL-APMS enables faithful proteome-scale mapping of native and endogenous galectin-glycoprotein networks. This framework for investigating galectin-glycoprotein interactions in their native context opens new avenues of broad glycobiology discovery and pinpoints critical targets for downstream galectin-centered translational research.

## Materials and Methods

### Cell Lines, Cell Culture, and Cell Treatments

Jurkat cells were obtained from the American Type Culture Collection (ATCC). Jurkat cells were cultured in RPMI-1640 medium supplemented with 10% fetal bovine serum (FBS), 1% penicillin-streptomycin-glutamine (PSG), and 0.1% beta-mercaptoethanol (BME). All cultures were incubated at 37°C in a humidified atmosphere containing 5% CO₂. For SILAC experiments, Jurkat cells were labeled for two weeks in Arginine and Lysine-free RPMI-1640 supplemented with 10% dialyzed fetal bovine serum (dFBS), 1% PSG, 0.1% BME, and 0.1 mg/ml light arginine and lysine or heavy arginine (Arg 10) and lysine (Lys 8). For pre-treatment of Kifunensine (KIF) and Swainsonine (SW), Jurkat cells were cultured in RPMI-1640 containing 10 µM KIF or 500 nM SW for 96 hours prior to downstream processing.

### Recombinant Galectin-3 Binding and Crosslinking

For conventional flow cytometry and flow-FRET experiments, Jurkat cells were plated on 96-well U bottom plates (5 x 10^5^ cells/well) and washed twice with cold 1x PBS. Following washing, cells were resuspended in 30 µl of 1.25 µg/ml recombinant galectin-3 (rGAL-3; R&D Systems 8259-GA), mixed gently, and incubated on ice for 20 min. After rGAL-3 incubation, cells were immediately crosslinked (DSG, DSSO, or DSP) at indicated concentrations, mixed gently, and incubated on ice for 20 mins. For competitive inhibition experiments, crosslinking was performed in 50 mM β-lactose, 50 mM D-sucrose, or 10 mM Thiodigalactoside (TDG). After 20 minutes of crosslinker reaction time, cell suspensions were diluted with cold FACS buffer (1% BSA, 0.1% NaN₃, 1X PBS) followed by two additional FACS washes (200 µl each). To pellet cells, plates were centrifuged at 350 RCF, 4°C and were aspirated by benchtop vacuum. For his-tagged galectin-3 experiments, N-terminal his-tagged galectin-3 (ACRO Biosystems GA3-H5247) and C-terminal his-tagged galectin-3 (ACRO Biosystems GA3-H5129) were crosslinked and stained congruent with untagged galectin-3 experiments or stained with anti-his (abcam ab117512).

For LC-MS experiments, Jurkat cells (100 million cells per condition) were washed twice with cold 1X PBS. Following washing, cells were resuspended in 500 µl of 40 µg/ml rGAL-3 and incubated at 4°C with rotation for 20 minutes. After rGAL-3 incubation, Jurkat cells were immediately crosslinked by adding 2 mM DSSO, mixing gently, and incubating at 4°C with rotation for 20 minutes. After 20 minutes of crosslinker reaction time, DSSO was diluted with cold 1X PBS followed by two additional 1X PBS washes (1 ml each). After washing, cells were resuspended in 500 µl of 80 µg/ml biotinylated anti-GAL-3 IgG (BioLegend 125404) and incubated at 4°C with rotation for 20 minutes. DSSO crosslinking of biotinylated anti-GAL-3 IgG and subsequent washing was performed congruent with that of rGAL-3.

### Generation of MGAT1 Knockout Jurkat Cells

Jurkat cells were nucleofected with Cas9-sgRNA ribonucleoprotein complex targeting exon 2 of MGAT1 (NM_002406.4) using the AMAXA Nucleofector II system. Following transfection, cells were recovered in RPMI with 10% FBS for 3 days. The efficiency of MGAT1 knockout was assessed by lectin stain with L-PHA, E-PHA, and DSL followed by flow cytometry. The L-PHA negative population was sorted by Fluorescence Assisted Cell Sorting (FACS) and expanded as the MGAT1 knockout population.

### Streptavidin enrichment and LC-MS sample preparation

Crosslinked cell pellets were lysed in 8 M urea buffer containing 0.3 M NaCl, 50 mM sodium phosphate (pH 8.0), 0.5% IGEPAL CA-630, and 1% SDS. Lysates (1 mL per 20 million cells) were homogenized via needle shearing (18G, 21G, and 23G, 20 passes each), followed by two rounds of 20 s sonication (1s on, 1s off). Lysates were centrifuged (16,000 × g, 15 min), and supernatants were incubated overnight at 4°C with streptavidin agarose resin (ThermoFisher; 112.5 µL slurry per mL of lysate). After overnight streptavidin incubation, beads were washed with lysis buffer (five washes, 20x bead volume each), followed by five washes in 8 M urea wash buffer (1.22 M NaCl, 0.5% SDS, 125 mM Tris-HCl pH 8.0, 10% ethanol, 10% isopropanol). Beads were washed with 25 mM ammonium bicarbonate (one wash, 20x bead volume), resuspended in 5.3 M urea in 25 mM ABC, and reduced with 5 mM TCEP (37°C, 15 min), followed by alkylation with 10 mM iodoacetamide (RT, dark, 30 min). Beads were digested on-bead first with LysC (1 µg per 112.5 µL bead volume, 4 h at 37°C), then diluted to 1.5 M urea and digested with trypsin (2.5 µg per 112.5 µL, 14 h at 37°C). Supernatants were acidified with 0.3% TFA to pH 3, desalted using C18 cartridges (pre-conditioned with ACN, 50% ACN, 0.5% acetic acid, and 0.1% TFA), and eluted with 80% ACN. Eluates were dried via SpeedVac and stored at –80°C.

### Reverse Phase Liquid Chromatography and Mass Spectrometry

Peptide mixtures were analyzed by liquid chromatography-tandem mass spectrometry (LC-MS/MS) using a Thermo Scientific™ UltiMate™ 3000 UHPLC system coupled online to a Thermo Scientific™ Orbitrap Fusion Lumos™ Tribrid™ Mass Spectrometer via a nano-electrospray ion source. Peptides were separated by reverse-phase chromatography on a 50 cm x 75 μm I.D. EasySpray Acclaim® PepMap™ RSLC C18 analytical column (Thermo Fisher Scientific) using a linear 90-minute (non-SILAC) or 120-minute (SILAC) gradient from 4% to 22% acetonitrile in 0.1% formic acid (solvent A: 0.1% formic acid in water; solvent B: 0.1% formic acid in acetonitrile) at a flow rate of 300 nL/min. Full MS scans were acquired in the Orbitrap over a mass range of 375 to 1500 m/z at a resolution of 60,000 (at m/z 400), followed by data-dependent MS/MS scans of the top 15 most intense precursor ions using higher collision-induced dissociation (HCD with a normalized collision energy of 30% in the linear ion trap). Dynamic exclusion was enabled with a duration of 30 seconds. The AGC target was set to 4 x 10^5^ for MS1 and 6 x 10^6^ for MS2, with maximum injection times of 50 ms and 35 ms, respectively. SILAC analyses were performed in biological duplicate across two independent experiments to ensure reproducibility.

### LC-MS Data Analysis

Protein identification and quantification from LC-MS/MS data were performed using MaxQuant (version 1.6.0.16)^97^. Raw mass spectrometry files were searched against the Homo sapiens (UP000005640_9606) or Mus musculus (UP000000589_10090) proteomes. MS/MS spectra were filtered to retain a maximum of eight fragment peaks per 100 Da interval. The precursor mass tolerance was set to 20 ppm for the initial search and 4.5 ppm for the main search. False discovery rates (FDR) for both peptide spectral matches and protein identifications were controlled at 1%, using the razor peptide approach. Trypsin/P was specified as the digestion enzyme, allowing up to two missed cleavages, with no allowance for nonspecific cleavage. Carbamidomethylation of cysteine residues was defined as a fixed modification, while methionine oxidation and N-terminal acetylation were set as variable modifications, allowing up to two variable modifications per peptide. Protein identifications were filtered by known cell culture contaminants and bulk RNA-seq expressed genes (TPM ≥ 1). Protein intensities were calculated as the integrated peak area across the chromatographic elution profile. For isotopic clusters, intensities from all isotopic peaks were summed. Only peptides classified as unique plus razor were used for quantification to ensure protein-level accuracy. For SILAC analyses, data were searched with a multiplicity of two with Arg 10 and Lys 8 heavy isotopic amino acids selected. Enrichment was calculated as Log₂ of wild type to MGAT1^-/-^ intensities and compared via one-sample, two-tailed t-test against a null enrichment of 0. Enrichment for proteins identified exclusively in one labeling condition was imputed as: 1 + *MAXIMUM FINITE ENRICHMENT*.

### Crosslinked Peptide (XL-MS) Sample Preparation and Data Analysis

Prior to MS^n^ analysis, crosslinked peptide enrichment was performed on digested peptides by size exclusion chromatography (SEC). Peptides were reconstituted in SEC mobile phase (0.1% formic acid and 30% ACN). Peptides separated via a Superdex Peptide PC 3.2/30 column (300 x 3.2 mm) at a flow rate of 50 µL/min and UV absorbance was read at 215, 245, and 280 nm. Two minute fractions were collected, and fractions 24 and 26 were analyzed. After LC-MS^n^ analysis, raw LC MS/MS and LC MS^n^ data were converted to MGF files using ProteoWizard MSConvert (v. 3.0.10738) respectively. MS^3^ and MS/MS spectra were searched via Batch-Tag within a developmental version of Protein Prospector (v.5.19.1, University of California, San Francisco) against a randomly concatenated decoy-containing *Homo sapiens* database (SwissProt.2017.11.01.random.concat; 20,240 entries). The mass tolerances for parent ions were set to ± 20 ppm and fragment ions were set to 0.6 Da. Trypsin was set as the enzyme with three maximum missed cleavages for MS^3^ data and two maximum missed cleavages for MS/MS data. Carbamidomethylation was set as a constant modification for cysteine, while protein N-terminal acetylation, N-terminal conversion of glutamine to pyroglutamic acid, asparagine deamidation, and methionine oxidation were selected as variable modifications. Three additional modifications were added for uncleaved lysines and protein N-termini: alkene (C3H2O, +54.0106 Da), unsaturated thiol (C3H2SO, +86.0041 Da), sulfenic acid (C3H4SO2, +104.0152 Da), corresponding to remnant moieties of DSSO after cross-link cleavage. The in-house software XL-Tools was used to automatically identify, summarize and validate cross-linked peptides based on Protein Prospector database search results and MS^n^ data. For MS/MS data, peptides were identified at 5% FDR for benchmarking analysis. For all MS3 analyses, the FDR of cross-linked peptide identifications was determined to be < 1%, based on previous reports^37^.

### Bioinformatic Analyses

Mouse thymocyte single cell RNA-sequencing data were sourced from Pankaew et al. 2022^42^. Consensus primary sequence, N-glycosylation site, extracellular region, protein repeat domain, and functional family annotation data were sourced from Uniprot. Cell surfaceome protein lists were sourced from Sánchez et al. 2026, Bausch Fluck et al. 2018 (SURFY), Diaz-Ramos et al. 2011, da Cunha et al. 2009, and Uniprot annotation of N-glycosylation^47–50^. Gene Ontology (BP and MF) and pathway (KEGG and Reactome) enrichment analyses were performed as described using the gProfiler engine against the Homo sapiens or Mus musculus proteomes with a user threshold of 0.05. Non-galectin-3 interaction data were sourced from StringDB physical subnetworks with a confidence threshold of 0.5. BioGRID, IntAct, StringDB data were sourced from the BioGRID (version 5.0.260), IntAct, and StringDB repositories with gene identifier LGALS3^52–55^.

### SDS-PAGE, Immunoblotting

Immunoblot sample preparations, lysis, and streptavidin enrichment were performed congruent with LC-MS preparations described above. Whole cell lysate sample protein concentrations were quantified by BCA protein quantification kit (Thermo Fisher Scientific). For each lane, LDS sample buffer (Thermo Fisher Scientific) was added to whole cell lysate, flow-through, or bead-bound proteins to produce a 1X working dilution. For DSP crosslinking experiments, LDS sample buffer was supplemented with 100 mM DTT to ensure complete reduction during boiling. Liquid samples were boiled for 5 mins. Bead-bound samples were boiled for 15 minutes to elute bound proteins. Proteins were resolved by SDS-PAGE using 1 mm 4-12% Bis-Tris gradient gels (Thermo Fisher Scientific) and transferred to Nitrocellulose membranes at 30V for 2 hours with ice submersion. Protein transfer was validated by Ponceau staining and membranes were blocked with 5% BSA in 1X Tris-Buffered Saline + Tween 20 for 1 hr at room temperature. Primary antibody and STREP-DYLIGHT-649 incubations were performed overnight at 4°C. Immunoblot images were acquired using the iBrightFL1500 system (Invitrogen) with smart exposure. To assess glycosylation status of target proteins, streptavidin-bound proteins were treated with Endo H or PNGase F (New England Biolabs) according to manufacturer’s protocols.

### MALDI-ToF Glycomics

For glycomics experiments, xGL-APMS preparations followed the LC-MS galectin-3 crosslinking methodology described above. Surfaceomics preparations were performed as previously described in Sánchez et al., 2026. Briefly, Jurkat cells were labeled via sulfo-NHS-biotin (Thermo Fisher Scientific). Cells were washed twice with PBS supplemented with 0.5% FBS, then resuspended in supplemented PBS at 1 x 10^8^ cells/ml. Sulfo-NHS-biotin was added at 0.5 mM and incubated at 4°C for 20 min. Reactions were quenched by adding 50 mM Tris-HCl (pH 8.0) and incubated for 5 min at 4°C. Cells were then washed twice in PBS and processed for streptavidin enrichment.

Enrichment samples were analyzed at the UCSD GlycoAnalytics core facility. N-glycans were released from glycoproteins using PNGase-F (New England Biolabs), followed by purification of the N-glycan by C18 and PGC cartridges. N-glycans were permethylated (PM) and further purified using C18 cartridge, finally the PM glycans were mixed with Super-DHB matrix and spotted on MALDI plate. Mass spectra were acquired on Bruker AutoFlex MALDI-ToF/ToF instruments in positive-reflectron mode. The spectral data was analyzed using GlycoWork Bench software and possible structures were annotated on the spectra. Glycomics analyses were performed in independent biological triplicate.

### CD5 Construct Generation and Expression

Wild type and glycan-deficient HA-CD5 constructs were designed based on the consensus human CD5 coding sequence from the UCSC genome browser (CCDS8000.1). A construct comprising an ATG start codon followed by an HA tag, a GSSSS linker, and the CD5 sequence was cloned into the pMax vector. Plasmid cloning and maxi-prep were performed by GenScript. Jurkat cell transfection was performed using the AMAXA Nucleofector II system and round cap 0.2 cm nucleofection cuvettes (USA Scientific) and nucleofection was initiated using a preset program (C-016). Nucleofected cells were incubated overnight in complete RPMI-1640 media prior to downstream processing. Expression was validated by surface anti-HA staining and conventional flow cytometry and imaging flow cytometry. Expected glycosylation states of each construct were confirmed by endoglycosidase treatment and immunoblotting.

### Panel Selection and Antibodies

Antibodies and recombinant lectins for immunophenotyping experiments were selected based on previous reports. For Jurkat validation experiments, xGL-APMS interactors and non-interactors were selected based on commercial availability of PE-conjugated formats. Non-interactors were selected as Jurkat surfaceome proteins that did not meet significant enrichment criteria. Non-interactors with untargeted surfaceomic intensity below minimum for xGL-APMS interactors were not considered for validation experiments. Proteins with non-detectable signal above isotype controls were not considered for validation experiments.

5NTE (AD2: PE, BioLegend, 344002: unconjugated, BioLegend, 344003), ADAM10 (SHM14: unconjugated, BioLegend, 352702), ITGB2 for flow cytometry (1B4/CD18: PE, BioLegend, 373408), ITGB2 for western blot (D4N5Z: unconjugated, Cell Signaling Technology, 73663S), CD1C (L161: PE, BioLegend, 331506: unconjugated, BioLegend, 331501), CD2 (RPA-2.10: PE, BioLegend, 300207: unconjugated, BioLegend, 300202), CD48 (BJ40: PE, BioLegend, 336708), CD5 (L17F12: PE, BioLegend, 364013: unconjugated, BioLegend, 364001), CD55 (JS11: unconjugated, BioLegend, 311301), CD59 (p282 [H19]: PE, BioLegend, 304707), CD6 (BL-CD6: PE, BioLegend, 313904: FITC, BioLegend, 313906), CD69 (FN50: PE, BioLegend, 310905: unconjugated, BioLegend, 310902), CD82 (ASL-24: PE, BioLegend, 342104), DPP10 (OTI1A5: unconjugated, Invitrogen, 503585S), HLA-E (3D12: PE, BioLegend, 342603), ICAM2 (CBR-IC2/2: PE, BioLegend, 328506: unconjugated, BioLegend, 328502), ICAM3 (CBR-IC3/1: PE, BioLegend, 330005), IGF2R (QA19A18: PE, BioLegend, 364204), IL10RB (S17009F: PE, BioLegend, 396803: unconjugated, BioLegend, 396802), IL2RG (QA17A55: PE, BioLegend, 314603: unconjugated, BioLegend, 314602), ITGA3 (ASC-1: PE, BioLegend, 343803), ITGA4 (9F10: PE, BioLegend, 304304), ITGAL (HI111: PE, BioLegend, 301202), ITGAV (NKI-M9: PE, BioLegend, 327910), ITGB1 (TS2/16: PE, BioLegend, 303004), LGALS3 for flow cytometry and immunoprecipitation (M3/38: PE, BioLegend, 125405: APC, BioLegend, 125420: biotin, BioLegend, 125404), LGALS3 for western blot (B2C10: unconjugated, BD Biosciences, BDB556904), PLXNA1 (8A10A6: unconjugated, Invitrogen, MA5-31856), PTPRC (human) for flow cytometry (H130: PE, BioLegend, 304002), PTPRC (human) for western blot (D9M8I: unconjugated, Cell Signaling Technology, 13917S), PTPRC (m) (QA17A26: PE, BioLegend, 157604), PTPRJ (MEM-CD148/05: PE, Invitrogen, A15799: unconjugated, Invitrogen, MA5-28592), SCARB1 (m1B9: PE, BioLegend, 363204), SLAMF6/NTB-A (NT-7: PE, BioLegend, 317208: unconjugated, BioLegend, 317202), TFRC (D7G9X: unconjugated, Cell Signaling Technology, 13113S), THY1 for FRET (S22015F: PE, BioLegend, 389906), THY1 for immunofluorescence (AF-9: unconjugated, abcam, ab23894), TNFRSF8 (BY88: FITC, BioLegend, 333902), TSPAN7 (B2D: FITC, Invitrogen, MA562253), ITGA6 (GoH3: PE, BioLegend, 313612), mouse IgG (H+L) secondary (polyclonal: Alexa Fluor 488, Invitrogen, A-21202: Alexa Fluor 647, Invitrogen, A-31571), HA.11 epitope tag (16B12: unconjugated, BioLegend, 901501), His tag (AD1.1.10: FITC, abcam, ab117512), CD4 (m) (RM4-5: PerCP-Cy5.5, BioLegend, 100540), CD8 (m) (53-6.7: PE-Cy7, BioLegend, 100722), CD19 (m) (6D5: Brilliant Violet 421, BioLegend, 115538), CD44 (m) (IM7: APC, BioLegend, 103012), CD25 (m) (PC61: PE, BioLegend, 102035), CD62L (m) (MEL-14: Brilliant Violet 421, BioLegend, 104436), and CD3 (m) (17A2: Brilliant Violet 711, BioLegend, 100241).

### Flow Cytometry, Flow-FRET and Imaging Flow Cytometry

Flow cytometry data were acquired using an Agilent NovoCyte Quanteon flow cytometer. Plated cells were blocked with FACS buffer and stained with primary antibodies in 96-well U bottom plates (50 µl/well) for 30 minutes according to the antibody manufacturer’s recommended per cell concentrations. Following staining, cell suspensions were diluted with cold FACS buffer (150 µl/well) followed by two additional washes with cold FACS buffer (200 µl/well). Stained cells were resuspended in 1% PFA prior to flow cytometry data acquisition.

Flow-FRET analyses were performed using potential glycoprotein ligands as FRET donors (PE conjugated IgG) and galectin-3 as FRET acceptor (APC-conjugated IgG) based on established antibody-based FRET methodology^59–61^. Briefly, crosslinked Jurkat cells were (i) single stained with PE-conjugated anti-glycoprotein, (ii) single stained with APC-conjugated anti-GAL-3, or (iii) co-stained with PE-anti-glycoprotein and APC-anti-GAL-3. Data were collected first using excitation by the blue (488 nm) laser with detection at 528/20 nm (donor intensity) and 667/30 nm (bleed/FRET), then using excitation by the red (637 nm) laser to directly quantify APC-anti-GAL-3 fluorescence. Donor-derived and acceptor-derived spectral bleed-through into 667/30 nm were calculated from regression models using single stained controls. Expected 667/30 nm signal was calculated per cell as the sum of donor-derived and acceptor-derived bleed-through. Per cell FRET was calculated as the difference between the observed and expected 667/30 nm signal. Normalized FRET was calculated as: 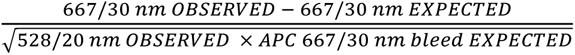, where 528/20 *nm OBSERVED* represents donor intensity and *APC* 667/30 *nm bleed EXPECTED* represents acceptor intensity.

For non-capping imaging flow cytometry experiments, Jurkat cells were plated on 96-well U bottom plates (0.5 x 10^6^ cells/well) with rGAL-3 incubation, crosslinking and staining as described above. Cells were additionally stained with 5 µg/ml Hoechst and 20 µg/ml anti-mouse Alexa Fluor (AF)-488. For capping assays, Jurkat cells were plated on 96-well U bottom plates (0.5 x 10^6^ cells/well) and washed twice with cold 1x PBS (200 µl/well). Following washing, cells were resuspended in 30 µl of 1.25 µg/ml recombinant human galectin-3 (rGAL-3), mixed gently, and incubated at 37°C for 30 mins. Following rGAL-3 incubation, APC-conjugated anti-GAL-3 IgG was immediately added at 2.5 µg/ml and incubated at 37°C for 20 mins to induce capping. Capped cells were DSG fixed and stained with primary mouse anti-glycoprotein IgG, Hoechst and anti-mouse AF-488. Flow cytometry images (n = 2500 per condition) were acquired on live focused cells (Hoechst^LOW^, Gradient RMS^HIGH^) cells using the AMNIS ImageStream®X Mark II system using the 60X objective at minimum flow rate. Co-localization was calculated using the *Similarity* feature in the AMNIS IDEAS analysis software using bright-field-derived cell membrane mask via the *Erode* function. Image display properties were set per-condition based on image pixel data with equal minimum thresholding across all conditions.

### Confocal Microscopy

Jurkat cells were plated on 96-well U bottom plates (0.5 x 10^6^ cells/well) with rGAL-3 incubation, crosslinking and staining as described above. Following staining, poly-lysine coated coverslips were washed twice with PBS and once with ddH2O, then mounted onto glass slides using Mowiol 4-88 mounting medium and allowed to cure overnight at room temperature. Confocal images were acquired using a Zeiss LSM 900 microscope equipped with Airyscan 2 detection and a 10x and 63x /1.4 NA oil-immersion objectives. Alexa-Fluor 647 was excited with the 640-nm laser line, Hoechst with the 405-nm laser line, and PE with the 568-nm laser line. Emission was collected using optimized Airyscan detector settings with 2.06 µs dwell time and 2x sample averaging. Z-slices were taken in 1 µm intervals with manually set first and last Z-positioning. Files were processed using ZEN Blue (v3.5.093, https://www.zeiss.com) using default settings. All imaging was performed under identical acquisition parameters between experimental conditions. Images were analyzed using Fiji (ImageJ, https://imagej.net/software/fiji). Co-localization was calculated per z-slice and per cell using the JaCoP plugin in Fiji with identical thresholding across conditions. Image display properties were set per-condition based on image pixel data with equal minimum thresholding across all conditions.

### Mice and Primary Cell Crosslinking

Wild-type C57BL/6J (000664) and galectin-3-deficient B6.Cg-*Lgals3^tm1Poi^*/J (006338) knockout mice were obtained from The Jackson Laboratory and maintained as previously described. Mice were housed at the University of California, Irvine in an environment-controlled, pathogen-free barrier facility on a 12-h/12-h light/dark cycle at constant temperature and humidity, with food and water available ad libitum. All mice were approved by the Institutional Animal Care and Use Committee (IACUC) of the University of California, Irvine (protocol # AUP-025-25). Mice were selected at random within designated experimental conditions and euthanized by CO2 and cervical dislocation prior to tissue extraction. For primary tissue crosslinking, harvested tissues were placed in 24 well plates, immediately homogenized by mashing in 1X PBS with or without 2 mM crosslinker and incubated on ice for 20 mins. Following crosslinking, cells were isolated via 70 µm mesh straining and processed for flow cytometry or confocal microscopy as described for Jurkat cells. For endogenous LC-MS experiments, cells from five thymuses were pooled for each genotype following lysis (5 mL lysis buffer), enrichment on streptavidin agarose (225 µl bead slurry), and LC-MS sample preparation as described above.

## Acknowledgements

The authors wish to thank Michael Hou of the University of California, Irvine Institute for Immunology Flow Cytometry core facility for support in flow-assisted cell sorting and imaging flow cytometry instrumentation, Adeela Sayed of the University of California, Irvine Optical Biology Core for microscopy instrumentation, and Biswa Choudury of the University of California, San Diego GlycoAnalytics Core for glycomics data acquisition and analysis. We also wish to thank Professor Michael Demetriou for useful discussion and critical feedback on the manuscript.

## CRediT authorship contribution statement

**Blake W Krisko**: Writing: review & editing, Writing: original draft, Visualization, Methodology, Investigation, Formal analysis.

**Aswath Balakrishnan**: Writing: review & editing, Visualization, Methodology, Investigation, Formal analysis.

**Andrew Wheat**: Writing: review & editing, Methodology, Investigation, Formal analysis, Data curation.

**Christofer Sánchez**: Writing: review & editing, Investigation.

**Clinton Yu**: Writing: review & editing, Data curation.

**Ria Bhakta**: Writing: review & editing, Investigation.

**Luna Witchey**: Writing: review & editing, Investigation.

**Minas Minasyan**: Writing: review & editing, Investigation.

**Lan Huang**: Writing: review & editing, Supervision, Funding acquisition.

**Haik Mkhikian**: Writing: review & editing, Writing: original draft, Visualization, Supervision, Methodology, Investigation, Funding acquisition, Formal analysis, Conceptualization.

---

**SUPPLEMENTARY FIGURE 1:**
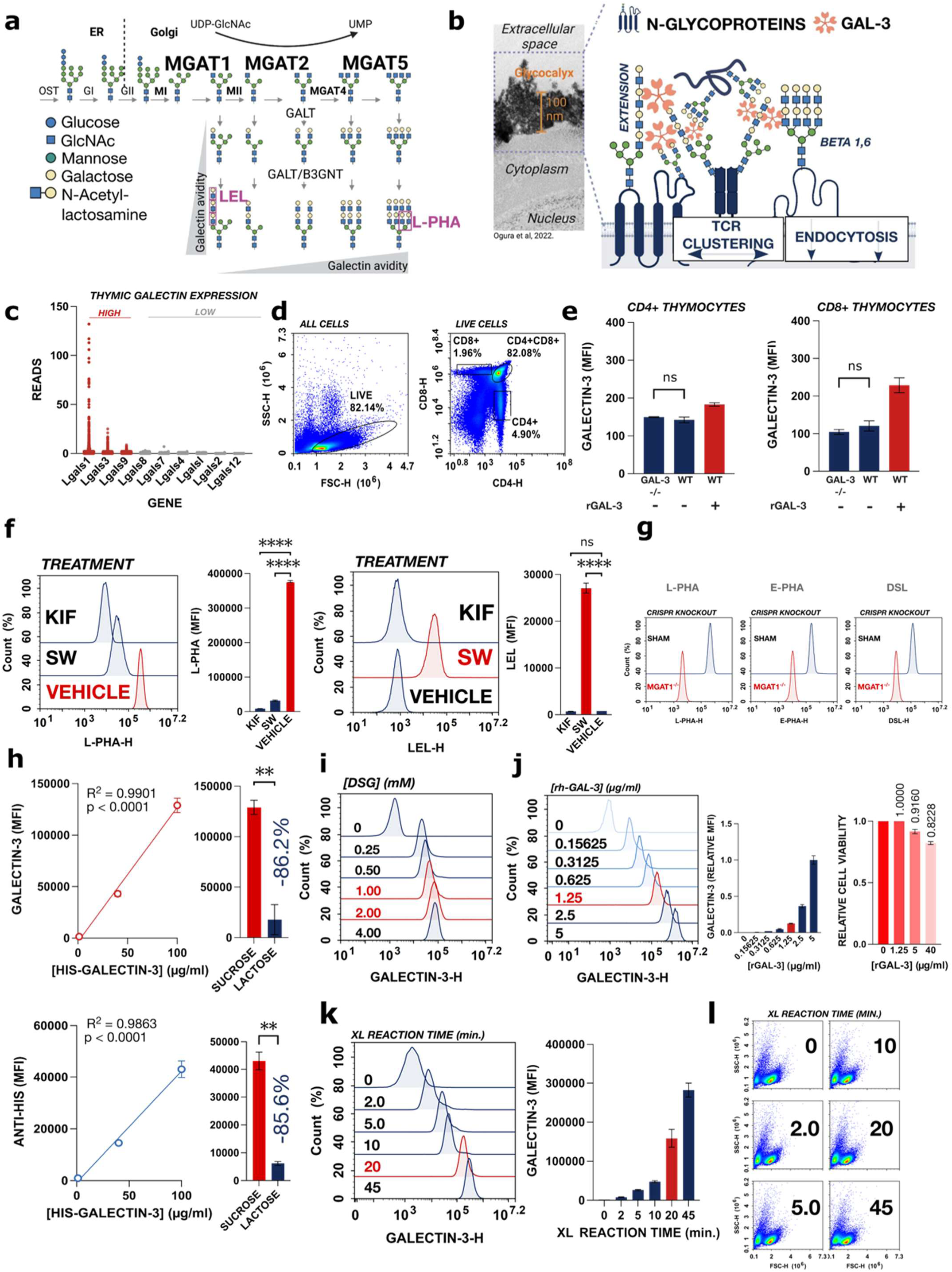
**a** Schematic of N-glycan branching (NGB) biosynthesis pathway. Golgi-resident N-acetylglycosaminyltransferases MGAT1, MGAT2, MGAT4 a/b, and MGAT5 add N-acetylglucosamine (GlcNAc) branches which are then variably extended by β-linked N-acetyllactosamine units. Binding preferences of plant lectins L-PHA and LEL, routinely used to quantify surface N-glycoforms, are indicated (pink). **b** Electron micrograph of Glycocalyx from Ogura et al. 2022 and schematic of galectin-3 lattice and associated T-cell functions. **c** Single cell RNA sequencing (scRNA-seq) of galectin family members from mouse thymocytes derived from Pankaew et al. 2022. **d** Representative flow cytometry gating strategy for CD4+ SP, CD8+ SP, and DP mouse thymocytes. **e** Quantification of surface galectin-3 on CD4+ SP and CD8+ SP thymocytes by routine flow cytometry. **f** Flow cytometry of L-PHA binding to Jurkat cells pre-treated with KIF and SW. **g** Flow cytometry of L-PHA, E-PHA, and DSL on wild type and MGAT1^-/-^ Jurkat cells. **h** Flow cytometry of titration of N-terminal his-tagged (NT-HIS) galectin-3 with crosslinking detected by anti-GAL-3 and anti-HIS IgG with corresponding sucrose vs. lactose controls using 100 µg/ml NT-HIS galectin-3. **i** Flow cytometry of galectin-3 following DSG crosslinker titration. **j** Flow cytometry of rGAL-3 titration on Jurkat cells with crosslinking and associated trypan blue viability staining following incubation with rGAL-3 alone. **k-l** Flow cytometry of crosslinking time course on Jurkat cells with 1.25 µg/ml rGAL-3 with corresponding light scattering characteristics (FSC-H/SSC-H). Flow cytometry experiments **(d-e)** were stained in triplicate. Flow cytometry experiments **(f-l)** were performed in biological triplicates. Data for **(d-l)** are representative of ≥ 3 independent experiments. Statistical comparisons by ordinary one-way ANOVA with Dunnett’s post-hoc test **(e-g)** and Welch’s t-test **(h)** (** = p ≤ 0.01, **** = p ≤ 0.0001).

**SUPPLEMENTARY FIGURE 2:**
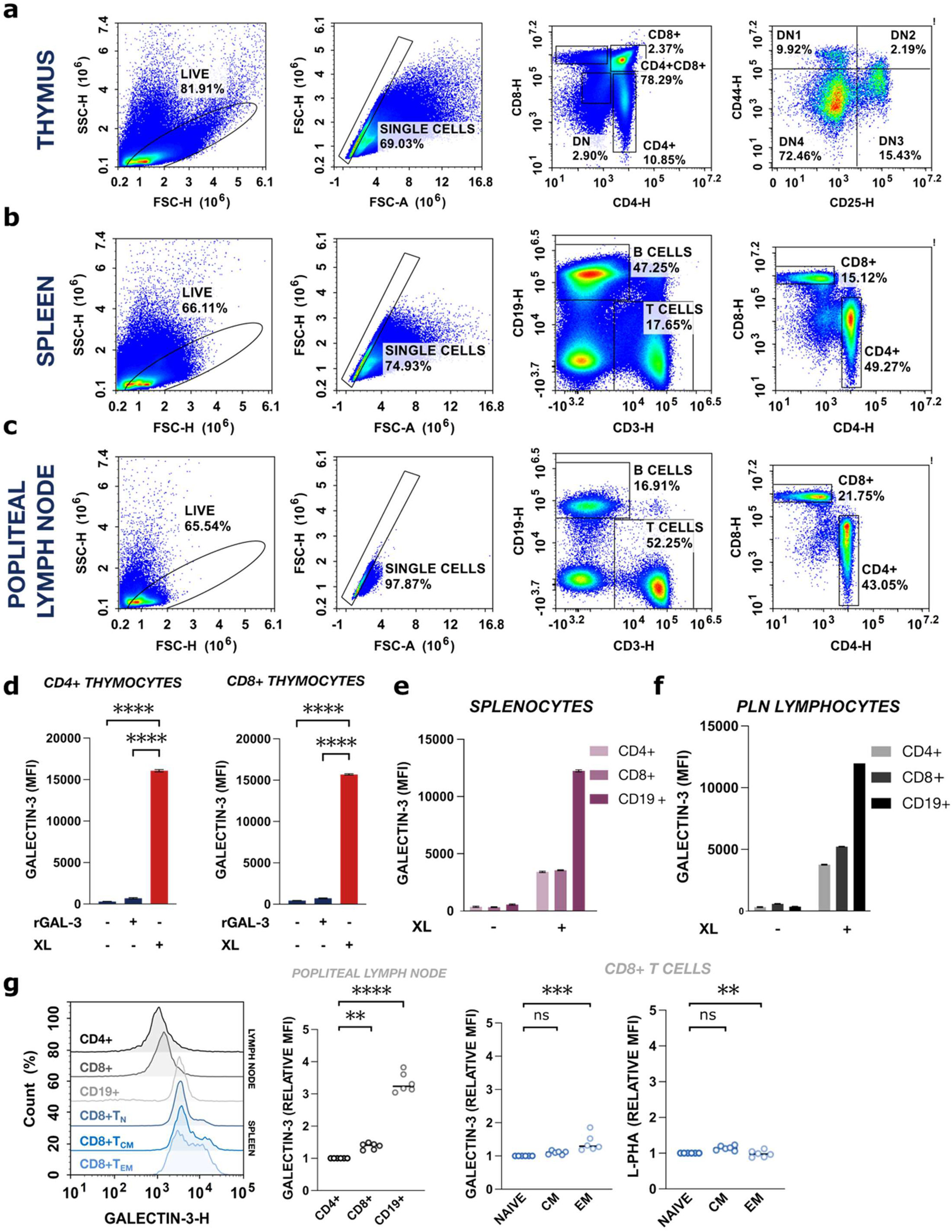
**a-c** Representative flow cytometry gating strategies for quantification of endogenous galectin-3 on thymocytes (a) (CD4 vs. CD8, CD25 vs. CD44) and peripheral immune cells (b-c) (CD3 vs. CD19, CD4 vs. CD8) following immediate crosslinking. **d** Flow cytometry of endogenous galectin-3 on CD4+ SP and CD8+ SP thymocytes under routine flow cytometry processing (non-crosslinked), incubation of rGAL-3 (non-crosslinked), and crosslinking conditions. **e-f** Flow cytometry of endogenous galectin-3 on splenic (e) and popliteal lymph node (PLN) (f) immune cells under non-crosslinked flow cytometry processing and crosslinking conditions. **g** Flow cytometry of endogenous galectin-3 on popliteal lymph node immune cells and naive, central memory, and effector memory CD8+ splenic T-cells with corresponding L-PHA binding. Flow cytometry experiments **(a-g)** were stained in triplicate. Data for **(a-g)** are representative of ≥ 3 independent experiments. Flow cytometry experiments were performed on age-matched wild type C57BL/6 mice (n = 4F, 2M, age = 7 weeks ± 2). Statistical comparisons by ordinary one-way ANOVA with Dunnett’s post-hoc test **(d, g)** (** = p ≤ 0.01, **** = p ≤ 0.0001).

**SUPPLEMENTARY FIGURE 3:**
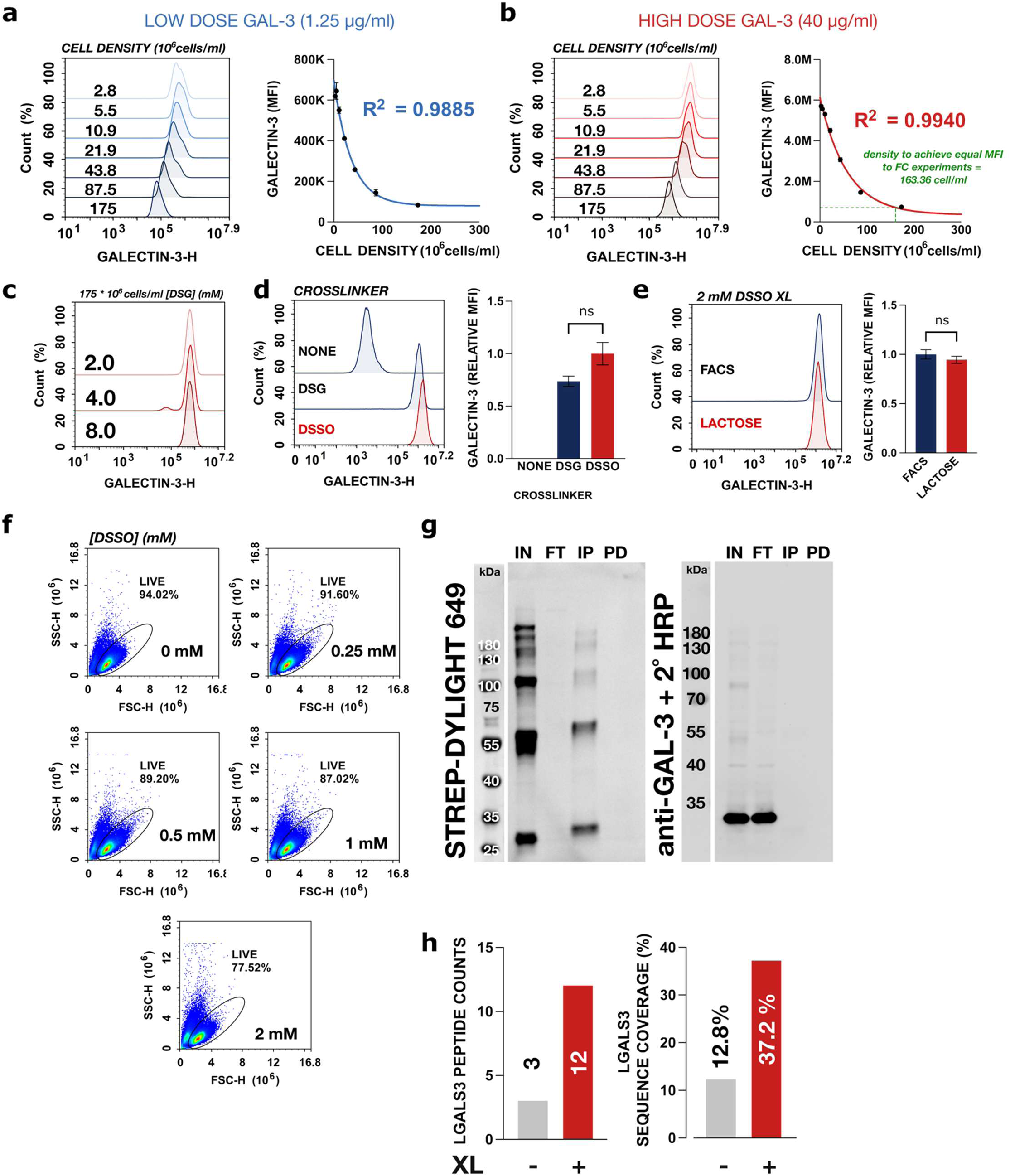
**a** Flow cytometry of galectin-3 on wild type Jurkat cells plated across a series of cell densities, with incubation of low dose (1.25 µg/ml) galectin-3 and DSG crosslinking. **b** Flow cytometry of galectin-3 on wild type Jurkat cells plated across a series of cell densities, with incubation of high dose (40 µg/ml) galectin-3 and DSG crosslinking. **c** Flow cytometry of galectin-3 following incubation of high dose galectin-3 (40 µg/ml) at high cell density (175 * 10⁶ cells/ml) across an extended titration of [DSG]. **d** Flow cytometry of galectin-3 following DSG crosslinking, DSSO crosslinking, and non-crosslinking controls. **e** Flow cytometry of galectin-3 following washing with 50 mM β-lactose vs. standard (FACS) washing buffer. **f** Light scattering (FSC-H vs. SSC-H) characteristics of Jurkat cells processed under the xGL-APMS workflow for DSSO titration immunoblotting. **g** Representative STREP-DYLIGHT-649 and anti-GAL-3 IgG + anti-mouse HRP of whole cell lysate (IN), flow-through (FT), and purified (IP) fractions of wild type and MGAT1^-/-^ Jurkat cells processed under non-crosslinked workflows. **h** LC-MS comparison of galectin-3 (LGALS3) peptide counts and sequence coverage with and without crosslinking. Data for **a-f** are representative of ≥ 3 independent experiments. Statistical comparisons by Welch’s t-test **(d-e)**.

**SUPPLEMENTARY FIGURE 4:**
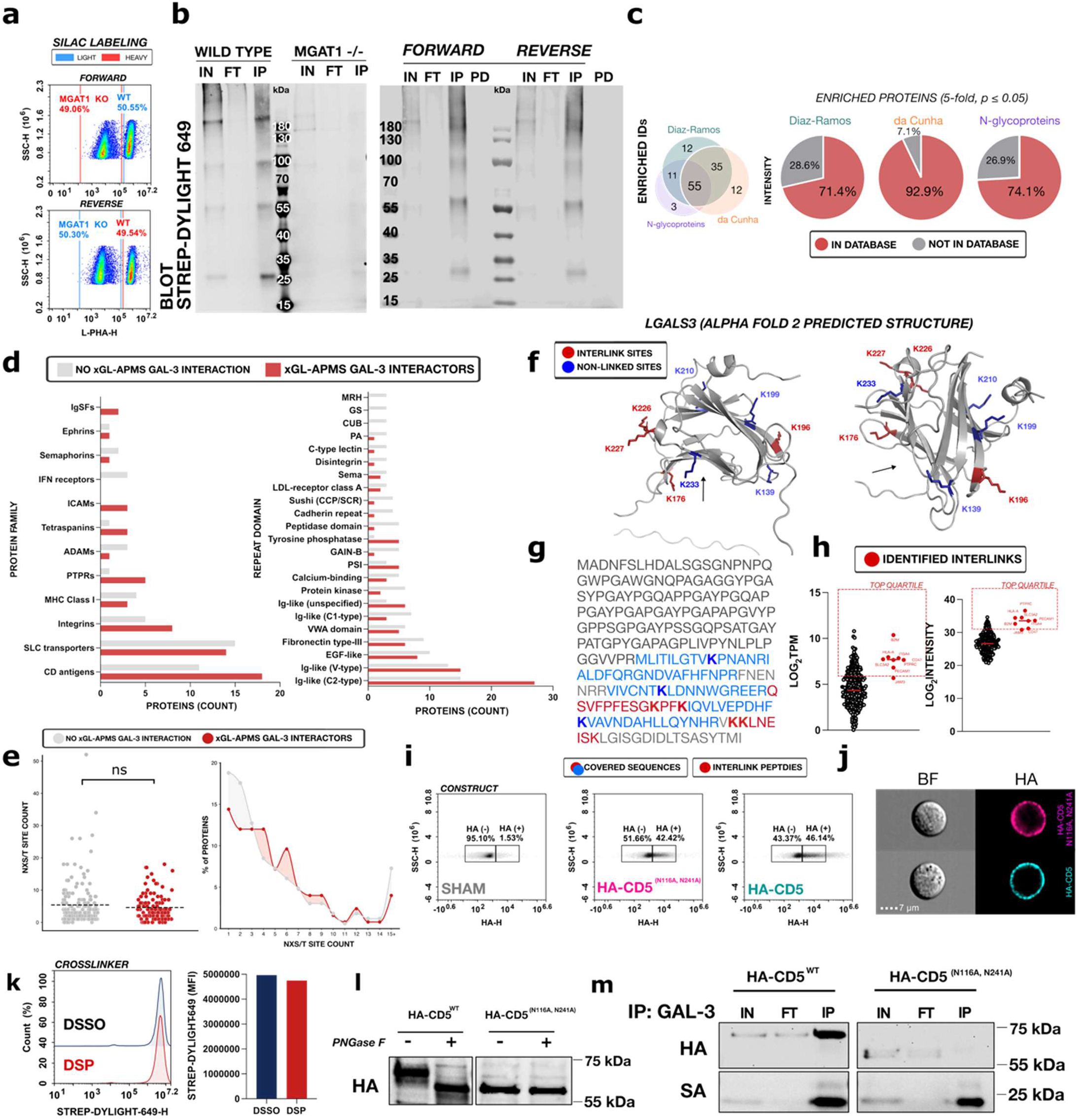
**a** Representative flow cytometry validation of 1:1 mixing of labeled wild type and MGAT1^-/-^ Jurkat cells for SILAC xGL-APMS experiments. **b** Representative STREP-DYLIGHT-649 immunoblot of whole cell lysate (IN), flow-through (FT), purified (IP), and post-digestion (PD) fractions of SILAC xGL-APMS experiments. **c** Venn diagram of significant hits from SILAC xGL-APMS experiments filtered by two surfaceome databases (Diaz-Ramos and da Cunha et al.) or by N-glycoproteins and Intensity share of significant hits from SILAC xGL-APMS experiments filtered by additional databases. **d** Protein family and repeat domain counts of the xGL-APMS galectin-3 interactome (red) and broader Jurkat surfaceome (grey). **e** Scatter dot plot of NXS/T site count on xGL-APMS-identified galectin-3 interactors (red) and broader surfaceome (grey) and relative share count within each group binned by NXS/T site number. **f** AlphaFold2 predicted structures of galectin-3 (LGALS3) with indicated crosslinked sites (red) and non-crosslinked sites (blue). Arrows indicate the carbohydrate binding pocket of galectin-3. **g** Sequence annotation of galectin-3 peptides covered in xGL-APMS experiments with corresponding crosslinked site annotations. **h** Bulk RNA sequencing (TPM) and surfaceomic intensity comparison of identified galectin-3 interlink partners (red) and broader Jurkat surface proteins (black). **i** Flow cytometry of anti-HA staining following SHAM transfection (grey), transfection with HA-CD5 (cyan), or transfection with HA-CD5^N116A,N241A^ (magenta). **j** Representative flow cytometry images of wild type Jurkat cells following transfection with HA-CD5 (cyan) or HA-CD5^N116A,N241A^ (magenta). **k** Flow cytometry of DSSO versus DSP galecitn-3 crosslinking. **l** Representative anti-HA immunoblot of whole cell lysates from wild type Jurkat cells transfected with HA-CD5 or HA-CD5^N116A,N241A^ and treated with the endoglycosidase PNGase F as indicated. **m** Representative anti-HA immunoblot of whole cell lysate (IN), flow-through (FT), and purified (IP) fractions following galectin-3 xGL-APMS preparation on wild type Jurkat cells transfected with HA-CD5 or HA-CD5^N116A,N241A^. Data from (**a, c-d, f, h)** are derived from four independent SILAC xGL-APMS experiments. Data for **(i-m)** are representative of ≥ 3 independent experiments.

**SUPPLEMENTARY FIGURE 5:**
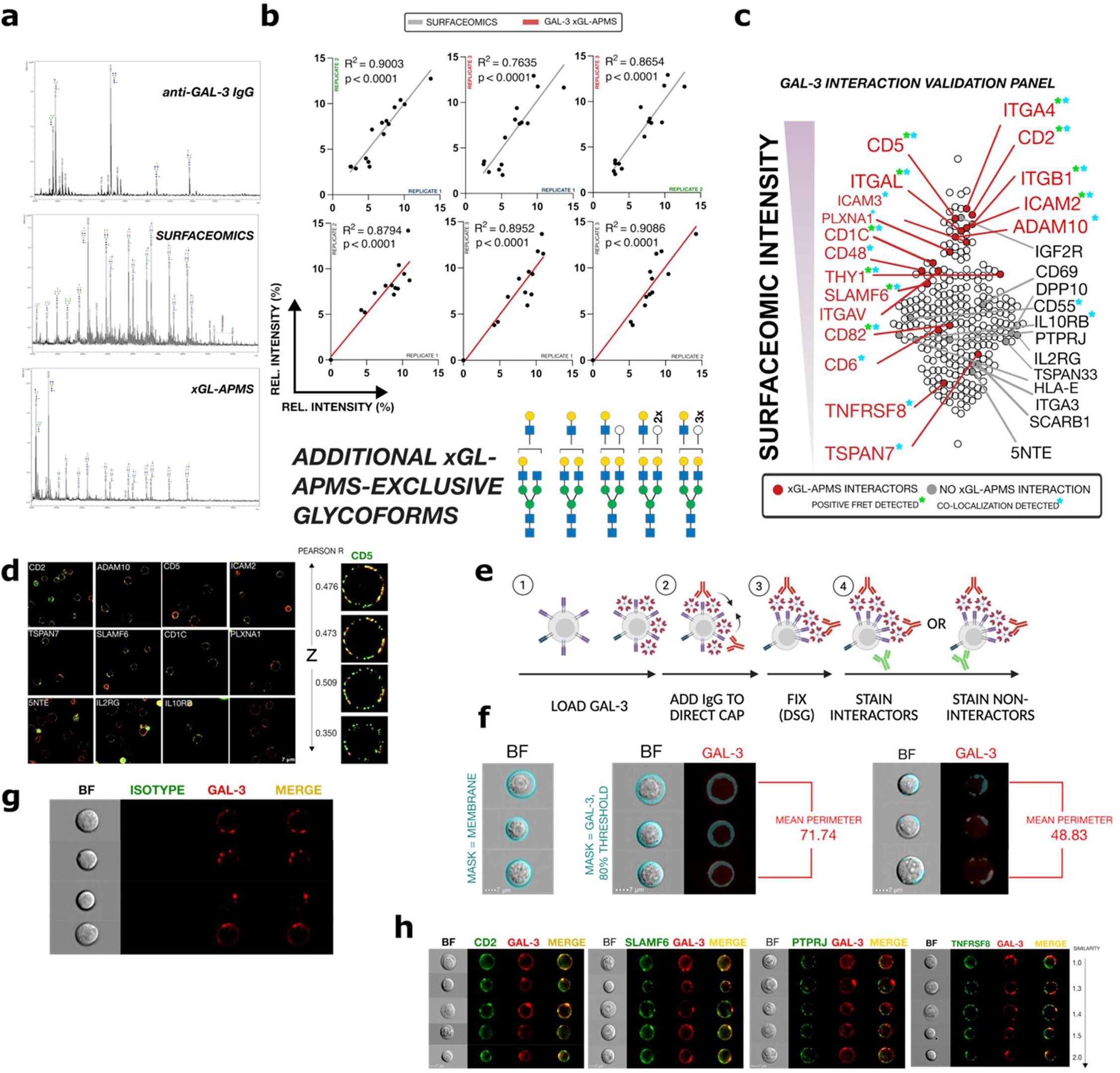
**a** Representative MALDI-ToF glycomic spectra of biotinylated anti-GAL-3 IgG alone, surfaceomics, and xGL-APMS preparations. N-glycoforms annotated in the anti-GAL-3 IgG isolate were not considered for quantitative analyses. **b** Linear regression of surfaceomics and xGL-APMS MALDI-ToF glycomics biological triplicates and N-glycoforms identified exclusively in one or two xGL-APMS MALDI-ToF glycomics replicates. **c** Overview of Jurkat surface proteins validated by flow-FRET, flow cytometry imaging (capping), or confocal microscopy with xGL-APMS interaction (red) or lacking xGL-APMS interaction (grey) mapped by surfaceomic intensity. **d** Representative 63X objective fields for surface receptors validated by confocal microscopy and 1 µm Z-slices for Jurkat surface proteins with xGL-APMS interaction (ADAM10, right) and lacking xGL-APMS interaction (5NTE, left) with corresponding Pearson’s R correlation for each slice. **e-f** Workflow schematic for capping experiments. Incubation of anti-GAL-3 IgG after lattice formation directs galectin-3 signal into distinct patches (quantifiable by reduced *Perimeter*) prior to fixation and staining for surface glycoproteins. **g** Representative flow cytometry images (n = 2500) of Mouse IgGκ1 isotype control + anti-Mouse AF-488 binding under capping conditions. **h** Additional representative flow cytometry images for surface proteins measured by capping. Data for **(e-h)** are representative of 3 independent experiments.

**SUPPLEMENTARY FIGURE 6:**
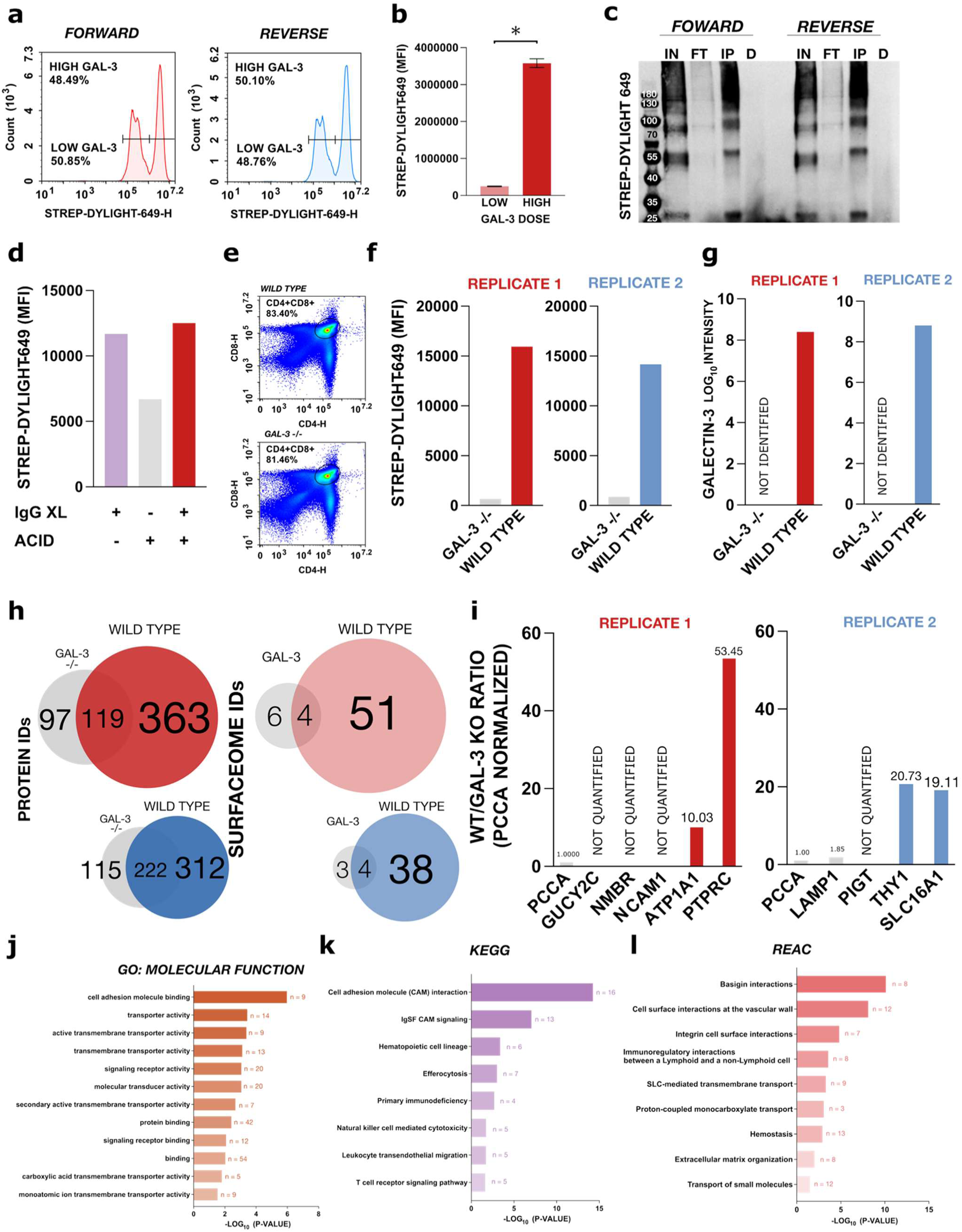
**a** Flow cytometry of STREP-DYLIGHT-649 step-down low dose galectin-3 and high dose galectin-3 SILAC xGL-APMS experiments confirming even mixing and quantitative gap. **b** Quantification of STREP-DYLIGHT-649 MFIs in mixed low dose GAL-3 and high dose GAL-3 SILAC xGL-APMS experiments (n = 2). **c** Representative STREP-DYLIGHT-649 immunoblot of whole cell lysate (IN), flow-through (FT), purified (IP), and post-digestion (PD) fractions from low dose galectin-3 and high dose galectin-3 SILAC xGL-APMS experiments. **d** Flow cytometry confirmation of anti-GAL-3 IgG crosslinking completeness by acid (150 mM NaCl, 20 mM HCl, pH 1.7) washing **e** Representative flow cytometry gating strategy for quantification of STREP-DYLIGHT-649 on wild type and Lgals3^-/-^ thymocytes confirming similar phenotype compositions for each genotype. **f** Flow cytometry quantification of STREP-DYLIGHT-649 signal on wild type and Lgals3^-/-^ CD4+/CD8+ thymocytes from endogenous xGL-APMS experiments. **g** LC-MS intensities of galectin-3 from endogenous xGL-APMS experiments. **h** Overlap of total protein IDs and surfaceome IDs from wild type and Lgals3^-/-^ thymocytes from endogenous xGL-APMS experiments. **i** Enrichment scoring of surfaceome proteins identified from both wild type and GAL-3 -/- xGL-APMS experiments. Raw intensity values were normalized to the endogenously biotinylated enzyme PCCA to correct for run-to-run variability. Proteins enriched ≥ 5-fold in wild type experiments were considered positive galectin-3 interaction hits. **j-l** Top GO: Molecular Function **(j),** KEGG **(k),** and REAC **(l)** enriched terms in the endogenous galectin-3 interactome. Statistical comparisons by Welch’s t-test **(b)** (* = p < 0.05)

## References

1. Hebert, D. N., Lamriben, L., Powers, E. T. & Kelly, J. W. The intrinsic and extrinsic effects of N-linked glycans on glycoproteostasis. Nat. Chem. Biol. 10, 902–910 (2014).

2. Davis, M. M. et al. Dynamics of Cell Surface Molecules During T Cell Recognition. Annu. Rev. Biochem. 72, 717–742 (2003).

3. Mkhikian, H., et al. *N* -Glycan Branching Regulates BTLA Opposite to PD-1 to Limit T Cell Hyperactivity Induced by Branching Deficiency. J. Immunol. 213, 1329–1337 (2024).

4. Mizutamari, R. K. et al. Reprogramming T cell fate through antibody-mediated galectin-1 blockade. in Methods in Cell Biology vol. 209 105–121 (Elsevier, 2026).

5. Yang, G.-S. et al. Galectin-3-Induced Glycan Lattices as Biophysical Modulators of Membrane Phase Behavior in Live Cells. ACS Chem. Biol. 21, 431–438 (2026).

6. Hu, X. et al. A comprehensive landscape of human organ N-glycoproteome. Nat. Commun. 17, 7656 (2026).

7. Dennis, J. W., Nabi, I. R. & Demetriou, M. Metabolism, cell surface organization, and disease. Cell 139, 1229–1241 (2009).

8. Schachter, H. The ‘yellow brick road’ to branched complex *N* -glycans. Glycobiology 1, 453– 461 (1991).

9. Kornfeld, R. & Kornfeld, S. ASSEMBLY OF ASPARAGINE-LINKED OLIGOSACCHARIDES. Annu. Rev. Biochem. 54, 631–664 (1985).

10. Grigorian, A., Torossian, S. & Demetriou, M. T-cell growth, cell surface organization, and the galectin-glycoprotein lattice. Immunol. Rev. 230, 232–246 (2009).

11. Yang, E. H. et al. Galectin-3 alters the lateral mobility and clustering of β1-integrin receptors. PLOS ONE 12, e0184378 (2017).

12. Demetriou, M., Granovsky, M., Quaggin, S. & Dennis, J. W. Negative regulation of T-cell activation and autoimmunity by Mgat5 N-glycosylation. Nature 409, 733–739 (2001).

13. Hirabayashi, J. et al. Oligosaccharide specificity of galectins: a search by frontal affinity chromatography. Biochim. Biophys. Acta 1572, 232–254 (2002).

14. Gilson, R. C., Gunasinghe, S. D., Johannes, L. & Gaus, K. Galectin-3 modulation of T-cell activation: mechanisms of membrane remodelling. Prog. Lipid Res. 76, 101010 (2019).

15. Xie, L. et al. Galectin-3 exacerbates autoimmune diabetes by limiting regulatory T cell differentiation and function. Sci. Adv. 12, eadz7916 (2026).

16. Lurje, I., Biegel, D., Zhang, I. & Tacke, F. The Role of Galectin-3 in Liver Inflammation and Fibrosis. J. Inflamm. Res. **Volume** 19, 1–21 (2026).

17. Hara, A. et al. Galectin-3 as a Next-Generation Biomarker for Detecting Early Stage of Various Diseases. Biomolecules 10, 389 (2020).

18. Lau, K. S. & Dennis, J. W. N-Glycans in cancer progression. Glycobiology 18, 750–760 (2008).

19. Mkhikian, H. et al. Age-associated impairment of T cell immunity is linked to sex-dimorphic elevation of N-glycan branching. *Nat*. Aging 2, 231–242 (2022).

20. Zhou, R. W. et al. Safe immunosuppression-resistant pan-cancer immunotherapeutics by velcro-like density-dependent targeting of tumor-associated carbohydrate antigens. Cell 188, 6737–6753.e18 (2025).

21. Mkhikian, H. et al. Genetics and the environment converge to dysregulate N-glycosylation in multiple sclerosis. Nat. Commun. 2, 334 (2011).

22. Chang, V. et al. Mapping galectin-3 ligands in human tear fluid establishes spliceoform-dependent lacritin binding. *Commun*. Biol. 10.1038/s42003-026-10337-2 (2026) doi:10.1038/s42003-026-10337-2.

23. Obermann, J. et al. Proteome-wide Identification of Glycosylation-dependent Interactors of Galectin-1 and Galectin-3 on Mesenchymal Retinal Pigment Epithelial (RPE) Cells. Mol. Cell. Proteomics MCP 16, 1528–1546 (2017).

24. Joeh, E. et al. Mapping glycan-mediated galectin-3 interactions by live cell proximity labeling. Proc. Natl. Acad. Sci. 117, 27329–27338 (2020).

25. Reeves, A. E., et al. Mapping the placental galectin-3 interactome identifies CD9 and ITGB1 as functional glycoprotein counterreceptors during syncytialization. Proc. Natl. Acad. Sci. 122, e2511042122 (2025).

26. Hwang, J. & Espenshade, P. J. Proximity-dependent biotin labelling in yeast using the engineered ascorbate peroxidase APEX2. Biochem. J. 473, 2463–2469 (2016).

27. Frankenfield, A. M., Fernandopulle, M. S., Hasan, S., Ward, M. E. & Hao, L. Development and Comparative Evaluation of Endolysosomal Proximity Labeling-Based Proteomic Methods in Human iPSC-Derived Neurons. Anal. Chem. 92, 15437–15444 (2020).

28. Zeng, Y., Ramya, T. N. C., Dirksen, A., Dawson, P. E. & Paulson, J. C. High-efficiency labeling of sialylated glycoproteins on living cells. Nat. Methods 6, 207–209 (2009).

29. Xie, Y. et al. Glycan–protein cross-linking mass spectrometry reveals sialic acid-mediated protein networks on cell surfaces. Chem. Sci. 12, 8767–8777 (2021).

30. Chen, S. et al. Quantitative Glycan-Protein Cross-Linking Mass Spectrometry Using Enrichable Linkers Reveals Extensive Glycan-Mediated Protein Interaction Networks. Anal. Chem. 97, 1584–1593 (2025).

31. Collas, P. Chop it, ChIP it, check it: the current status of chromatin immunoprecipitation. Front. Biosci. 13, 929 (2008).

32. Collas, P. The State-of-the-Art of Chromatin Immunoprecipitation. in Chromatin Immunoprecipitation Assays (ed. Collas, P.) vol. 567 1–25 (Humana Press, Totowa, NJ, 2009).

33. Guerrero, C., Tagwerker, C., Kaiser, P. & Huang, L. An Integrated Mass Spectrometry-based Proteomic Approach. Mol. Cell. Proteomics 5, 366–378 (2006).

34. Kaake, R. M. et al. A New in Vivo Cross-linking Mass Spectrometry Platform to Define Protein–Protein Interactions in Living Cells. Mol. Cell. Proteomics 13, 3533–3543 (2014).

35. Kandur, W. V., Kao, A., Vellucci, D., Huang, L. & Rychnovsky, S. D. Design of CID-cleavable protein cross-linkers: identical mass modifications for simpler sequence analysis. Org. Biomol. Chem. 13, 9793–9807 (2015).

36. Wang, X. & Huang, L. Dissecting Dynamic and Heterogeneous Proteasome Complexes Using In Vivo Cross-Linking-Assisted Affinity Purification and Mass Spectrometry. in The Ubiquitin Proteasome System (eds Mayor, T. & Kleiger, G.) vol. 1844 401–410 (Springer New York, New York, NY, 2018).

37. Wheat, A., et al. Protein interaction landscapes revealed by advanced in vivo cross-linking– mass spectrometry. Proc. Natl. Acad. Sci. 118, e2023360118 (2021).

38. Jiao, F. et al. Two-Dimensional Fractionation Method for Proteome-Wide Cross-Linking Mass Spectrometry Analysis. Anal. Chem. 94, 4236–4242 (2022).

39. Fasci, D., Van Ingen, H., Scheltema, R. A. & Heck, A. J. R. Histone Interaction Landscapes Visualized by Crosslinking Mass Spectrometry in Intact Cell Nuclei. Mol. Cell. Proteomics 17, 2018–2033 (2018).

40. Gutierrez, C. et al. Structural dynamics of the human COP9 signalosome revealed by cross-linking mass spectrometry and integrative modeling. Proc. Natl. Acad. Sci. 117, 4088–4098 (2020).

41. Zhou, R. W. et al. N-glycosylation bidirectionally extends the boundaries of thymocyte positive selection by decoupling Lck from Ca2+ signaling. Nat. Immunol. 15, 1038–1045 (2014).

42. Pankaew, S. et al. Multiplexed single-cell RNA-sequencing of mouse thymic and splenic samples. STAR Protoc. 3, 101041 (2022).

43. Clark, M. C. et al. Galectin-3 binds to CD45 on diffuse large B-cell lymphoma cells to regulate susceptibility to cell death. Blood 120, 4635–4644 (2012).

44. Chen, I.-J., Chen, H.-L. & Demetriou, M. Lateral Compartmentalization of T Cell Receptor Versus CD45 by Galectin-N-Glycan Binding and Microfilaments Coordinate Basal and Activation Signaling. J. Biol. Chem. 282, 35361–35372 (2007).

45. Mkhikian, H. et al. Golgi self-correction generates bioequivalent glycans to preserve cellular homeostasis. eLife 5, e14814 (2016).

46. Elbein, A. D., Tropea, J. E., Mitchell, M. & Kaushal, G. P. Kifunensine, a potent inhibitor of the glycoprotein processing mannosidase I. J. Biol. Chem. 265, 15599–15605 (1990).

47. Bausch-Fluck, D. et al. The in silico human surfaceome. Proc. Natl. Acad. Sci. 115, (2018).

48. Sánchez, C. D. et al. Improved T Cell Surfaceomics by Depleting Intracellularly Labelled Dead Cells. Mol. Cell. Proteomics MCP 25, 101503 (2026).

49. Díaz-Ramos, M. C., Engel, P. & Bastos, R. Towards a comprehensive human cell-surface immunome database. Immunol. Lett. 134, 183–187 (2011).

50. Da Cunha, J. P. C. et al. Bioinformatics construction of the human cell surfaceome. Proc. Natl. Acad. Sci. 106, 16752–16757 (2009).

51. Bernhard, O. K., Sheil, M. M. & Cunningham, A. L. Lateral Membrane Protein Associations of CD4 in Lymphoid Cells Detected by Cross-Linking and Mass Spectrometry. Biochemistry 43, 256–264 (2004).

52. Oughtred, R. et al. The BioGRID interaction database: 2019 update. Nucleic Acids Res. 47, D529–D541 (2019).

53. Oughtred, R. et al. The BIOGRID database: A comprehensive biomedical resource of curated protein, genetic, and chemical interactions. Protein Sci. 30, 187–200 (2021).

54. Szklarczyk, D. et al. The STRING database in 2023: protein–protein association networks and functional enrichment analyses for any sequenced genome of interest. Nucleic Acids Res. 51, D638–D646 (2023).

55. del Toro, N. et al. The IntAct database: efficient access to fine-grained molecular interaction data. Nucleic Acids Res. 50, D648–D653 (2022).

56. Plewka, J., Barnowski, K. & Rolak, L. GAL-3 protein with lactose: 28vp. 10.2210/pdb28vp/pdb (2026).

57. Wright, J. K., Tschopp, J. & Jaton, J. C. Preparation and characterization of chemically defined oligomers of rabbit immunoglobulin G molecules for the complement binding studies. Biochem. J. 187, 767–774 (1980).

58. Stanley, P. Golgi Glycosylation. Cold Spring Harb. Perspect. Biol. 3, a005199–a005199 (2011).

59. Kumar, A., Kremer, K. N., Sims, O. L. & Hedin, K. E. Measuring the proximity of T-lymphocyte CXCR4 and TCR by fluorescence resonance energy transfer (FRET). Methods Enzymol. 460, 379–397 (2009).

60. Kumar, A. et al. CXCR4 Physically Associates with the T Cell Receptor to Signal in T Cells. Immunity 25, 213–224 (2006).

61. Xia, Z. & Liu, Y. Reliable and Global Measurement of Fluorescence Resonance Energy Transfer Using Fluorescence Microscopes. Biophys. J. 81, 2395–2402 (2001).

62. Mezu-Ndubuisi, O. J. & Maheshwari, A. The role of integrins in inflammation and angiogenesis. Pediatr. Res. 89, 1619–1626 (2021).

63. Martz, E., Davignon, D., Kürzinger, K. & Springer, T. A. The Molecular Basis for Cytolytic T Lymphocyte Function: Analysis with Blocking Monoclonal Antibodies. in Mechanisms of Cell-Mediated Cytotoxicity (eds Clark, W. R. & Golstein, P.) vol. 146 447–468 (Springer US, Boston, MA, 1982).

64. Larson, R. S., Corbi, A. L., Berman, L. & Springer, T. Primary structure of the leukocyte function-associated molecule-1 alpha subunit: an integrin with an embedded domain defining a protein superfamily. J. Cell Biol. 108, 703–712 (1989).

65. Bourguignon, L. Y. W., Hyman, R., Trowbridge, I. & Singer, S. J. Participation of histocompatibility antigens in capping of molecularly independent cell surface components by their specific antibodies. Proc. Natl. Acad. Sci. 75, 2406–2410 (1978).

66. Bourguignon, L. Y. & Singer, S. J. Transmembrane interactions and the mechanism of capping of surface receptors by their specific ligands. Proc. Natl. Acad. Sci. 74, 5031–5035 (1977).

67. Huang, Q. et al. Imaging flow cytometry: from high - resolution morphological imaging to innovation in high - throughput multidimensional biomedical analysis. Front. Bioeng. Biotechnol. 13, 1580749 (2025).

68. Frankenfield, A. M. et al. Benchmarking SILAC Proteomics Workflows and Data Analysis Platforms. Mol. Cell. Proteomics 24, 100980 (2025).

69. Shi, S. et al. Inactivation of the Mgat1 gene in oocytes impairs oogenesis, but embryos lacking complex and hybrid N-glycans develop and implant. Mol. Cell. Biol. 24, 9920–9929 (2004).

70. Ioffe, E. & Stanley, P. Mice lacking N-acetylglucosaminyltransferase I activity die at mid-gestation, revealing an essential role for complex or hybrid N-linked carbohydrates. Proc. Natl. Acad. Sci. U. S. A. 91, 728–732 (1994).

71. Ioffe, E., Liu, Y. & Stanley, P. Complex *N* -glycans in *Mgat1* null preimplantation embryos arise from maternal *Mgat1* RNA. Glycobiology 7, 913–919 (1997).

72. Waas, M. et al. SurfaceGenie: a web-based application for prioritizing cell-type-specific marker candidates. Bioinformatics 36, 3447–3456 (2020).

73. Ramya, T. N. C. et al. In situ trans ligands of CD22 identified by glycan-protein photocross-linking-enabled proteomics. Mol. Cell. Proteomics MCP 9, 1339–1351 (2010).

74. Han, S., Collins, B. E., Bengtson, P. & Paulson, J. C. Homomultimeric complexes of CD22 in B cells revealed by protein-glycan cross-linking. Nat. Chem. Biol. 1, 93–97 (2005).

75. Tu, L. et al. L-selectin binds to P-selectin glycoprotein ligand-1 on leukocytes: interactions between the lectin, epidermal growth factor, and consensus repeat domains of the selectins determine ligand binding specificity. J. Immunol. 157, 3995–4004 (1996).

76. Xin, M., Dong, X.-W. & Guo, X.-L. Role of the interaction between galectin-3 and cell adhesion molecules in cancer metastasis. Biomed. Pharmacother. 69, 179–185 (2015).

77. Sedlář, A., et al. Interaction between Galectin-3 and Integrins Mediates Cell-Matrix Adhesion in Endothelial Cells and Mesenchymal Stem Cells. Int. J. Mol. Sci. 22, 5144 (2021).

78. Friedrichs, J., Manninen, A., Muller, D. J. & Helenius, J. Galectin-3 Regulates Integrin α2β1-mediated Adhesion to Collagen-I and -IV. J. Biol. Chem. 283, 32264–32272 (2008).

79. Matarrese, P. et al. Galectin-3 overexpression protects from apoptosis by improving cell adhesion properties. Int. J. Cancer 85, 545–554 (2000).

80. Carturan, A., et al. Overcoming the Lack of CD2:CD58 Costimulation in Chimeric Antigen Receptor T-Cell Immunotherapies. Blood 144, 504–504 (2024).

81. Liu, J., Chow, V. T. K. & Jois, S. D. S. A novel, rapid and sensitive heterotypic cell adhesion assay for CD2–CD58 interaction, and its application for testing inhibitory peptides. J. Immunol. Methods 291, 39–49 (2004).

82. Janssen, B. J. C. et al. Structural basis of semaphorin–plexin signalling. Nature 467, 1118– 1122 (2010).

83. Zhang, C. et al. SLC3A2 N-glycosylation and Golgi remodeling regulate SLC7A amino acid exchangers and stress mitigation. J. Biol. Chem. 299, 105416 (2023).

84. Murtazina, R., Booth, B. J., Bullis, B. L., Singh, D. N. & Fliegel, L. Functional analysis of polar amino-acid residues in membrane associated regions of the NHE1 isoform of the mammalian Na^+^ /H^+^ exchanger. Eur. J. Biochem. 268, 4674–4685 (2001).

85. Galić, S., Schneider, H.-P., Bröer, A., Deitmer, J. W. & Bröer, S. The loop between helix 4 and helix 5 in the monocarboxylate transporter MCT1 is important for substrate selection and protein stability. Biochem. J. 376, 413–422 (2003).

86. Nimmanon, T. et al. The ZIP6/ZIP10 heteromer is essential for the zinc-mediated trigger of mitosis. Cell. Mol. Life Sci. CMLS 78, 1781–1798 (2021).

87. Taylor, K. M. et al. Zinc transporter ZIP10 forms a heteromer with ZIP6 which regulates embryonic development and cell migration. Biochem. J. 473, 2531–2544 (2016).

88. Kouo, T. et al. Galectin-3 Shapes Antitumor Immune Responses by Suppressing CD8+ T Cells via LAG-3 and Inhibiting Expansion of Plasmacytoid Dendritic Cells. Cancer Immunol. Res. 3, 412–423 (2015).

89. Chatterjee, M., Kis-Toth, K., Thai, T.-H., Terhorst, C. & Tsokos, G. C. SLAMF6-driven co-stimulation of human peripheral T cells is defective in SLE T cells. Autoimmunity 44, 211– 218 (2011).

90. Na, K. et al. CD81 and CD82 expressing tumor-infiltrating lymphocytes in the NSCLC tumor microenvironment play a crucial role in T-cell activation and cytokine production. Front. Immunol. 15, 1336246 (2024).

91. Elishmereni, M. & Levi-Schaffer, F. CD48: A co-stimulatory receptor of immunity. Int. J. Biochem. Cell Biol. 43, 25–28 (2011).

92. Meyaard, L., Hurenkamp, J., Clevers, H., Lanier, L. L. & Phillips, J. H. Leukocyte-associated Ig-like receptor-1 functions as an inhibitory receptor on cytotoxic T cells. J. Immunol. 162, 5800–5804 (1999).

93. Dong, B. et al. CD5-mediated inhibition of TCR signaling proceeds normally in the absence of SHP-1. Int. J. Mol. Med. 38, 45–56 (2016).

94. Marie-Cardine, A. et al. SHP2-interacting Transmembrane Adaptor Protein (SIT), A Novel Disulfide-linked Dimer Regulating Human T Cell Activation. J. Exp. Med. 189, 1181–1194 (1999).

95. Sreejit, G. et al. The ESAT-6 Protein of Mycobacterium tuberculosis Interacts with Beta-2-Microglobulin (β2M) Affecting Antigen Presentation Function of Macrophage. PLoS Pathog. 10, e1004446 (2014).

96. Sagi, Y., Landrigan, A., Levy, R. & Levy, S. Complementary costimulation of human T-cell subpopulations by cluster of differentiation 28 (CD28) and CD81. Proc. Natl. Acad. Sci. 109, 1613–1618 (2012).

97. Cox, J. et al. Andromeda: A Peptide Search Engine Integrated into the MaxQuant Environment. J. Proteome Res. 10, 1794–1805 (2011).

